# Identification of negative BOLD responses using windkessel models

**DOI:** 10.1101/392290

**Authors:** Pedro A. Valdés-Hernández, Byron Bernal, Arash Moshkforoush, Catalina Dunoyer, Hui Ming Khoo, Jorge Bosch-Bayard, Nicolas von-Ellenrieder, Jean Gotman, Jorge J. Riera

## Abstract

Alongside positive BOLD responses (PBR), a variety of negative BOLD responses (NBR) with distinct underlying mechanisms also occur. We identify five mechanisms of NBR: **i**) local/lateral/contralateral inhibition (LCI), **ii**) neuronal disruption of network activity (NDA), **iii**) altered balance of neuro-metabolic/vascular couplings (ANC), **iv**) arterial blood stealing (ABS), and **v**) venous blood backpressure (VBB). Detecting and classifying these mechanisms from BOLD signals is pivotal in understanding normal/pathological brain functions. This requires models and parameters with anatomical/functional interpretation that furnish the understanding of how these mechanisms are fingerprinted by their BOLD responses. Here, we used a windkessel model with viscoelastic compliance as well as dynamics of both neuronal and tissue/blood O_2_ to investigate the generation, detection, classification and interpretation of the BOLD hemodynamic response functions (HRF) of above mechanisms. Firstly, we evaluated the use of the general linear model to detect simulated NBRs. Secondly, we tested the ability of a machine learning classifier, built from a simulated ensemble of HRFs, to predict the mechanism underlying a new HRF. Crossvalidation indicates NDA and ANC can accurately be classified solely from fMRI BOLD signals; while LCI, ABS and VBB might require additional imaging modalities. Thirdly, we demonstrated that estimators of the model parameters determinant in the NBRs formation are accurate, and precise to certain resolutions. Finally, we successfully applied our detection/classification/estimation methodology to EEG-fMRI data in a clinical situation where several of these mechanisms could coexist. We believe that the proper identification and interpretation of NBR mechanisms have important clinical and cognitive implications in fMRI studies.

## Introduction

It is well known that BOLD hemodynamic response functions (HRFs) exhibit a repertoire of shapes and polarities, reflecting a variety of underlying mechanisms determined by the interrelation between the excitability of neurons, cerebral blood flow (CBF), cerebral blood volume (CBV), cerebral metabolic rate of oxygen (CMRO_2_), oxygen extraction fraction (OEF) and blood deoxy-hemoglobin (dHb) content. The coupling between neuronal activity and positive BOLD responses (PBR) has been widely accepted in the literature (Buxton et al., 2004; Havlicek et al., 2017). On the contrary, there is a variety of experimental evidence attributing negative BOLD responses (NBRs) to multiple factors including vascular phenomena (Bandettini, 2012; Goense et al., 2012; Harel et al., 2002), neuronal inhibition (Maggioni et al., 2016; Pasley et al., 2007; Shmuel et al., 2002; Smith et al., 2004; Wade, 2002) or the combination of both (Shmuel et al., 2002). We believe the detection, classification, and interpretation of the underlying NBR mechanisms from BOLD signals is important to properly understand brain function in normal and pathological conditions.

In this paper, we identify/study/analyze five main NBR mechanisms:

i. Neuronal deactivation due to inhibition. This is the most accepted mechanism for NBR (Maggioni et al., 2016; Pasley et al., 2007; Schäfer et al., 2012; Shmuel et al., 2002; Smith et al., 2004; Wade, 2002). It accommodates any type of cortical and intracortical inhibition driven by afferents from an active region. The inhibition can be lateral, surrounding the PBR region (Shmuel et al., 2006; Wade, 2002), in nearly homotopic contralateral areas of the brain (Boorman et al., 2010; Schäfer et al., 2012; Stefanovic et al., 2004)—both seen during task-related fMRI experiments and electrical stimulation—among others involving cortico-thalamic-cortical loops. The inhibition can also occur in the same active region, but exceeding the excitation, provoking negative changes in the overall neuronal activity, resulting in an NBR. This is observed, for example, in epilepsy at the seizure onset zone (Pittau et al., 2013), where a significant slow EEG wave, presumably owing to a profound hyperpolarization of pyramidal cells (Neckelmann et al., 2000; Pollen, 1964), follows a fast interictal epileptic spike. For simplicity, we shall refer to these mechanisms as the *local/lateral/contralateral inhibition (LCI)*. We distinguish between *local* and *remote*, depending on whether the inhibition occurs in the active region or in lateral/contralateral regions, respectively.
ii. Deactivation (or disruption) of resting state networks (RSNs), such as the default mode network (DMN). This phenomenon has been observed during interictal epileptic discharges (IEDs) (Fahoum et al., 2013; Kobayashi et al., 2006; Laufs et al., 2007) as well as neuro-electrical wrist stimulation (Klingner et al., 2010). We shall refer to this mechanism as *neuronal disruption of network activity (NDA)*.
iii. Increase in CMRO_2_ without the expected activity-induced increase in relative CBF. This mismatch between blood supply and demand is provoked, mainly in regions affected by certain pathologies, by a disproportionate coupling between neural activity and CMRO_2_ (neurometabolic coupling), an insufficient coupling between neural activity and CBF (neurovascular coupling), or a combination of both mechanisms. For instance, NBR associated with an increase in neuronal activity was reported in the hippocampus of epileptic rats during seizures (Schridde et al., 2008) as well as in the ipsilateral somatosensory cortex of rats during electric forepaw stimulation (Devor et al., 2008). We shall refer to this mechanism as *altered balance between neuro-vascular/metabolic couplings (ANC)*.
iv. *Arterial blood stealing (ABS)*, in which CBF and CBV decrease in a region, thus exhibiting NBR, because its arterial supply is stealth by a nearby region with PBR that is demanding more blood flow. Therefore, this NBR is not directly related to changes in neuronal activity. This vascular phenomenon was firstly reported in the surroundings of the suprasylvian gyrus (areas 21a and 7) in the visual cortex of cats after visual stimulation, with concurrent PBR in V1 (areas 17 and 18) (Harel et al., 2002). Although not measured during the experiment, the authors suspected that no change in neuronal activity was expected to happen in the NBR region, according to direct measurements in experiments with similar stimulation paradigms. Stronger evidence in favor of the existence of this mechanism was recently provided by (Ma et al., 2017), also using visual stimulation of regions 17 and 18 of cats. This time, the lack of change in neuronal activity was confirmed using simultaneous local field potential (LFP) and multi-unit activity (MUA). More evidence was observed in rats during hindlimb electrical stimulation using simultaneous optical imaging and electrophysiological recordings (Hu and Huang, 2015).
v. *Venous blood backpressure (VBB)*, in which a region with NBR experiences a decrease in CBF but an unexpected increase in CBV (Goense et al., 2012), especially in the middle layers of the cortex. It is hypothesized that the NBR region receives an extra pressure from the venous compartment (Bandettini, 2012; Goense et al., 2012).

The nature and dynamics of these NBR mechanisms are different. Thus, it is reasonable to expect different associated HRF waveforms which could be used as fingerprints of the underlying mechanism. To verify and take advantage of these differences for the identification of the mechanisms, it is necessary to have biophysical models, beyond the usual model-free parameterization of (Glover, 1999). Such models allow one to understand how the dynamics of hidden physiological state variables and anatomical/functional parameters determine the observed BOLD responses. There is a paucity of models aiming to explain NBRs in the literature—two attempts to our knowledge. First, a detailed model of a local vascular anatomic network (VAN) predicted a decrease in CBF and CBV in branches surrounding an area undergoing a hyperemic response to a localized decrease in arteriole resistance (Boas et al., 2008). These CBF and CBV negativities can only have a pure vascular explanation owing to the vascular circuitry modeled in VAN. The authors also predicted an increase of dHb in these surrounding areas, which all together results in NBR. VAN is a highly detailed and comprehensive model that accounts for vessel resistance, compliance and oxygen delivery. Second, (Havlicek et al., 2017) addressed the possible role of the balance between inhibitory and excitatory activities on the formation of the PBRs and NBRs by adding two state neuronal equations to the extended balloon model (Friston, 2002): the 2S-DCM model. Authors fitted this model to the responses obtained by (Shmuel et al., 2006) in monkeys during visual stimulation, being able to reproduce the differences between the PBR and NBR.

However, including dynamics of neuronal activity, as well as its coupling to CMRO_2_ through tissue/blood O_2_ dynamics and to CBF through vasoactive signals, was beyond the scope of the VAN model; as it also was predicting BOLD responses. Besides, due to the complexity of the model, it is difficult to ascertain which parameters of VAN are determinant of the negativity in CBF and CBV. On the other hand, the 2S-DCM model does not explicitly account for the resistance of its elements and their interdependency, which is determined by vessel compliance under common intracranial blood pressure. This is necessary to model vascular circuitry. Hence, a common modeling framework, which expands on previous models, is needed to accommodate all of the above-mentioned NBR mechanisms.

In this paper, we propose to investigate the generation of NBRs using a more inclusive, though parsimonious, windkessel-based model with viscoelastic nonlinear delayed compliance (Zheng and Mayhew, 2009). To account for LCI and NDA, we propose to include two state (inhibitory and excitatory) equations, like (Havlicek et al., 2017), to describe the modulation of the dynamics of neuronal activity. To also account for ABS and VBB, we connect two windkessels, representing regions with PBR and NBR, by a common supplying artery and a common draining vein. The windkessel model with delayed compliance (Mandeville et al., 1999) allows explicit model of a circuit of nonlinear “blood resistors”. It also allows for the inclusion of nonlinear blood inertial forces using circuit-equivalent inductive elements (Spronck et al., 2012). In the windkessel framework, CBF modulation by vasoactive signals induced by neuronal activity (the neurovascular coupling) is proxied by a state equation for arteriole blood resistance. Moreover, for the modulation of CMRO_2_ by neuronal activity (the neurometabolic coupling) we import the state equations from the oxygen to tissue transport (OTT) model of (Zheng et al., 2002). These equations account for the dynamics of the OEF and the O_2_ concentrations in both tissue and blood. This incorporation is necessary to simulate the neuro-metabolic/vascular imbalance in ANC.

Using our model, we predict a family of HRFs for each of the above-mentioned NBR mechanisms and investigate if they are detectable and classifiable from fMRI data—to our knowledge, no study has addressed this in the literature. We are interested in the extreme cases of the model, i.e. particular values of its parameters in which the NBR mechanisms purely manifest. To that end, we identify those parameters, and range of values, that are determinant in the formation of the NBR of each mechanism during fast single and block inputs. These ranges of parameters map to an ensemble of BOLD HRFs for each mechanism which its distinguishability from the rest of the ensembles we aim to investigate. In order to obtain such ensembles, we perform trials of stochastic noisy realistic fMRI simulations of the particular sub-models with random values of their respective parameter ranges and estimate the associated HRFs. We subsequently build a machine learning ensemble classifier which uses the HRFs as features and their corresponding mechanisms as classes. We evaluate the performance of the classifier in predicting mechanisms from their BOLD signals. The presence of noise poses a question about the detectability of the HRFs from fMRI data. To investigate this, we test the ability of the general linear model (GLM) (Friston et al., 1995) to detect them. As stated above, one of the benefits of providing a biophysical model to explain the abovementioned NBRs is we can obtain anatomical and functional information of the underlying mechanism from its estimated parameters. Hence, we also test the accuracy and precision in their estimation.

Finally, we applied these methods to probe the presence of the NBR mechanisms in some cases of focal epilepsy, which is one of the current research interests in our lab. We chose this pathology because it has shown a variety of puzzling HRF waveforms during interictal epileptiform discharges (IEDs) (Rathakrishnan et al., 2010). We believe that classification and mechanistic interpretation could have important clinical implications.

## Materials and Methods

### Notation

A lower case bold symbol, e.g. ***x***, denotes a column vector. An upper case bold symbol, e.g. **X**, denotes a 2×2 matrix. Non-bold symbols, e.g. *x*, denote scalar variables. **1** and **0** are 2-lentgh column vectors of ones and zeros, respectively. **I**_*N*_ is the *N*×*N* identity matrix. The superscript *T* denotes transposition. ***x***∼*N*(***m,* Σ**) means ***x*** is multinormal distributed with mean ***m*** and covariance Σ. *In*(*x*) is the natural logarithm of *x* and 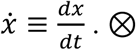. denotes temporal convolution. If 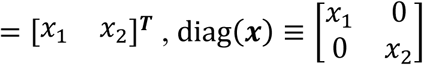.

### A biophysical model for NBR mechanisms

The model connects two windkessel regions (1 and 2), as depicted in Figure 1 and it has the following state-space form:

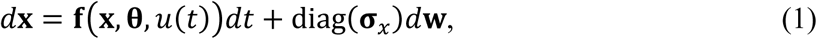

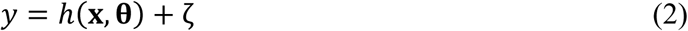

**Figure 1.**
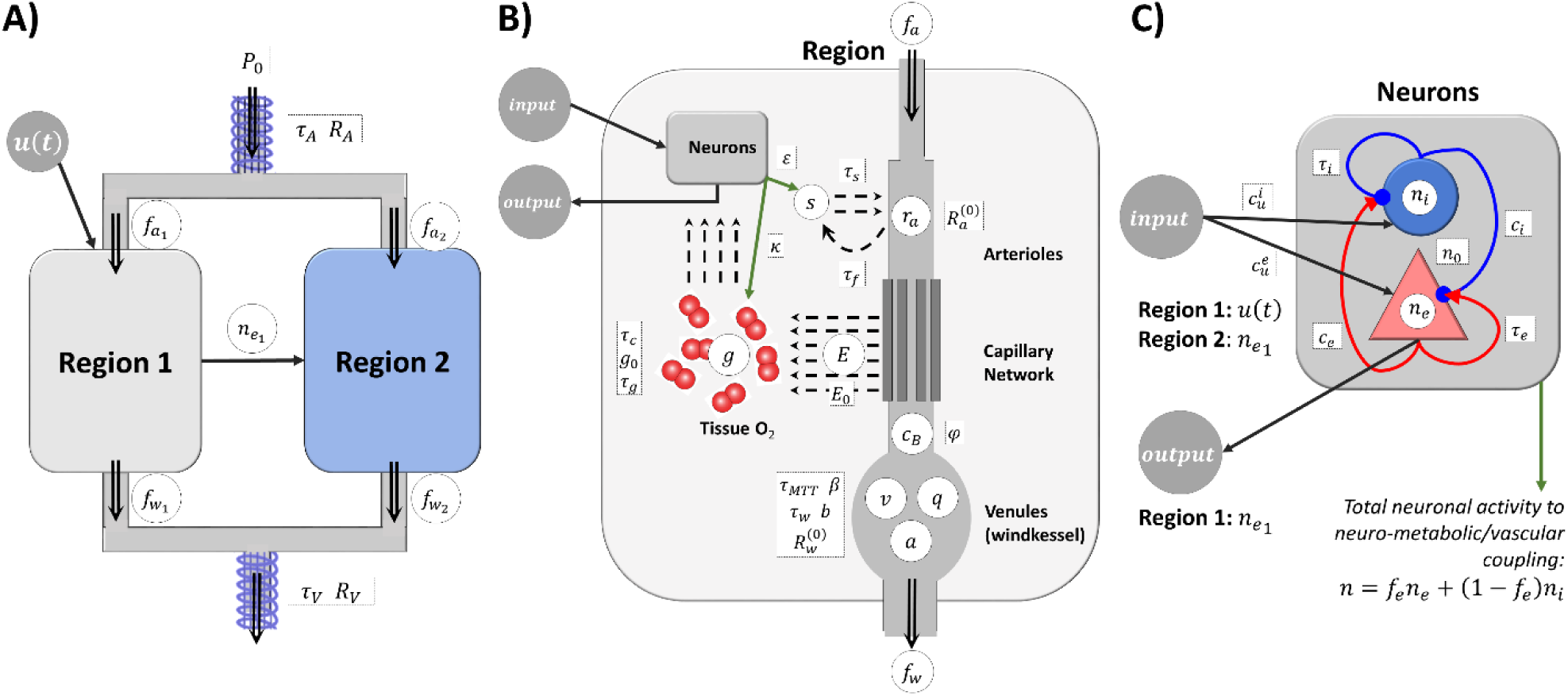
The model of connected windkessels with viscoelastic compliance, two-state neuronal dynamics and tissue/blood O_2_ dynamics. White circles represent the state variables outlined in Table 1. Symbols in the squares represent model parameters outlined in

where **X** is the vector of state variables, **θ** is the vector of model parameters, **f** is the drift function, **σ**_*x*_ is the vector of variances of state variables (the diffusion function), *h* the observation function, *d***w**∼*N*(**0**,**I**) is a Brownian stochastic process and *ζ∼N*(0,*σ*). The system is driven by an input *u*(*t*). The state variables, parameters and equations for both drift and observation terms are summarized in Table 1, Table 2 and Table 3, respectively.

Table 2. Double arrows represent CBF. Black bold labels denote different intra- and extra-cellular compartments relevant to the models. A) Regions 1 and 2 are connected by a common supplying artery and a common draining vein, both with blood resistance and inertia. The whole system is under a common constant pressure. The external input acts on region 1. The latter influences the dynamics of region 2 by means of neuronal or vascular. B) Model details each region. The input acts (continuous black arrows) on neuronal activity, which is represented by a black box with an input/output pair. Neurovascular and neurometabolic couplings are represented with green arrows. That is, changes in the total neuronal activity, i.e. the weighted sum of excitatory and inhibitory contributions, causes the release of a vasoactive signal that is autoregulated by CBF (neurovascular coupling). O_2_ is extracted by tissue and consumed by neurons. This extraction (OEF) depends on a set of state equations that model its dynamics, as well as those of tissue and blood O_2_ concentration. Neuronal activity also modulates the CMRO_2_ (neurometabolic coupling). These dynamics determine the amount of dHb in the windkessel. The windkessel volume is determined by the difference between the input and output CBF and viscoelastic properties. The BOLD signal is determined by the CBV and dHb, relative to their baseline values. C) Neuronal model. Two states—excitatory and inhibitory—are self- and influencing each other through effective connectivity. The input to region 1 is the external input, while the input to region 2 is the excitatory activity of region 1.

**Table 1.** States variables (and their steady state, baseline or initial values) of the biophysical models. A relative variable *x* is defined as the ratio of its value to its baseline, i.e. *X/X*^(0)^. These state variables are defined for each region. We omit the 1,2 subscripts for brevity.

| State variable | Steady state /initial value/ baseline |
| --- | --- |
| Relative excitatory neuronal activity ($n_e$) | 1 |
| Relative inhibitory neuronal activity ($n_i$) | 1 |
| Vasoactive signal ($s$) | 0 |
| Relative arteriole resistance ($r_a$) | 1 |
| Relative cerebral blood volume ($v$) | 1 |
| Relative cerebral blood flow in the arteriole ($f_a$) | 1 |
| Relative cerebral blood flow in the windkessel ($f_w$) | 1 |
| Relative pressure to volume relation constant ($a$) | 1 |
| Relative amount of deoxy-hemoglobin in the balloon/windkessel compartment ($q$) | 1 |
| Oxygen extraction fraction (OEF) ($E$) | $E_0$ |
| Tissue $O_2$ concentration normalized by plasma $O_2$ concentration in the arterial end of the capillary ($g$) | $G_0$ |
| Total blood $O_2$ concentration normalized by blood $O_2$ concentration in the arterial end of the capillary ($c_B$) | $G_0 - E_0 / \ln(1 - E_0(1 - G_0)^{-1})$ |

**Table 2.**
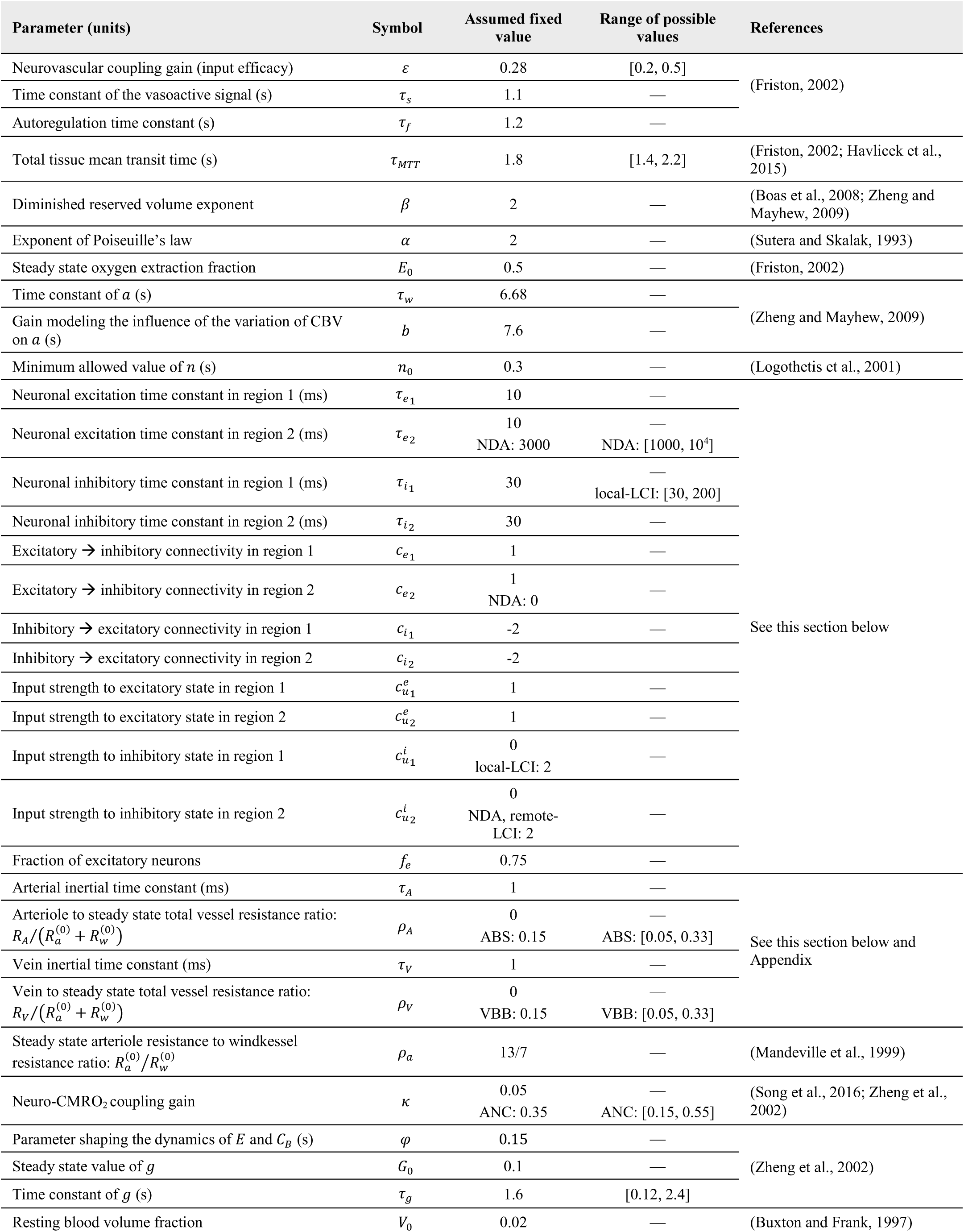
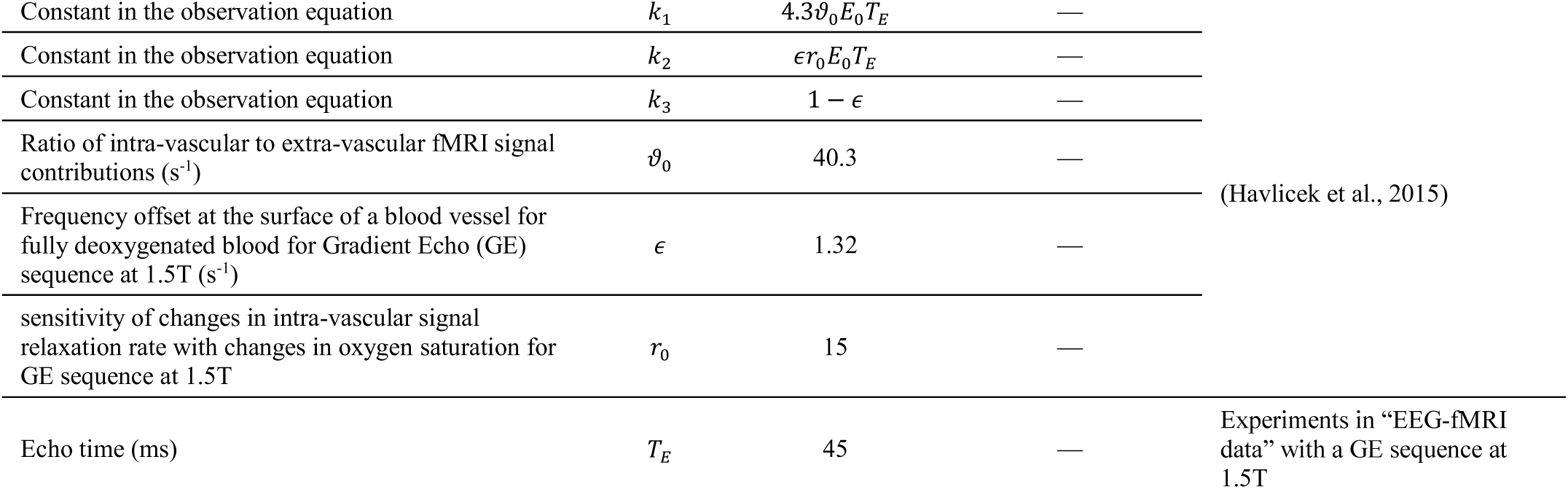
Parameters of the biophysical models. The assumed fixed values are those used in simulations performed to investigate the detectability of the NBR using GLM. In the estimation of the parameters, the values are allowed to vary within the provided ranges. These are also the ranges from which the parameters are randomly sampled to generate the ensemble of HRFs used to build the machine learning classifier. Note that these ranges are not the most general ranges, but those leading to NBR mechanisms. That is the case of *c*_*i*_, and 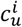, as explained in “Particular cases of the model”; *κ*, which yields PBR responses for values below 0.1s^−1^; and *τ*_*g*_ which also yields PBR responses for values higher than 6s. The values in this table are used for all models, unless specified otherwise. There is no need to provide uncertainty to all parameters, only a subset is enough to provide the entire possible span of variability in the HRFs.

**Table 3.**
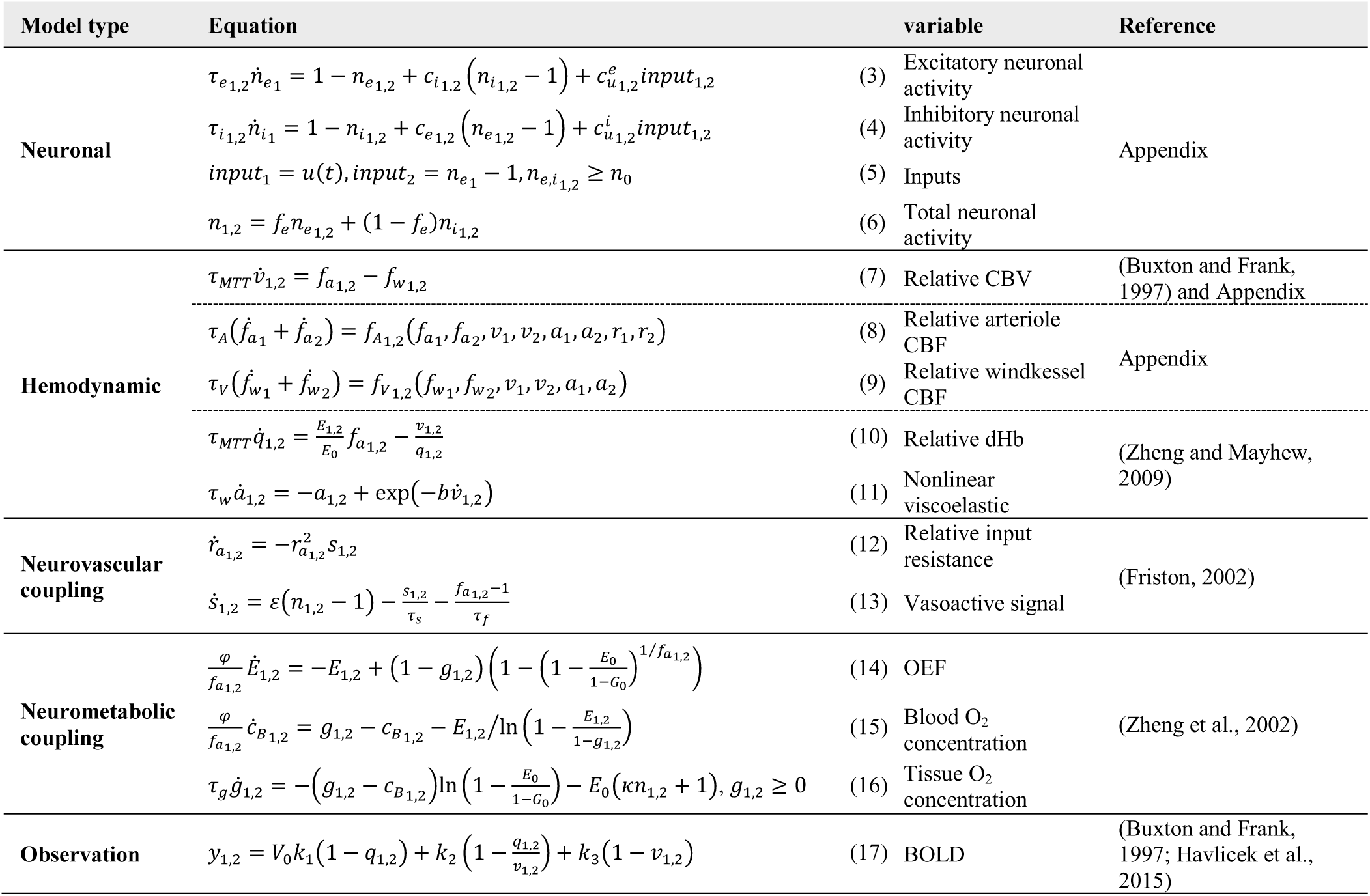
Equations of the drift term in the state equation in (1) for each region. The system is driven by an input *u(t)*.

### Particular cases of the model

To determine the extreme ensembles in the HRF space, corresponding to the situations in which each NBR mechanisms purely manifest, we consider particular cases of the model. These sub-models are used for simulation of the HRF ensembles and for investigating the detection of NBRs in fMRI experiment. For the sake of parsimony, we also use them for parameter estimation. The values of the parameters for all these particular situations are summarized in

### Table 2. Note that local-LCI and ANC are the only mechanism that do not need two interacting regions to happen, so we can disconnect them and consider a model with only one region

**Figure 2.**
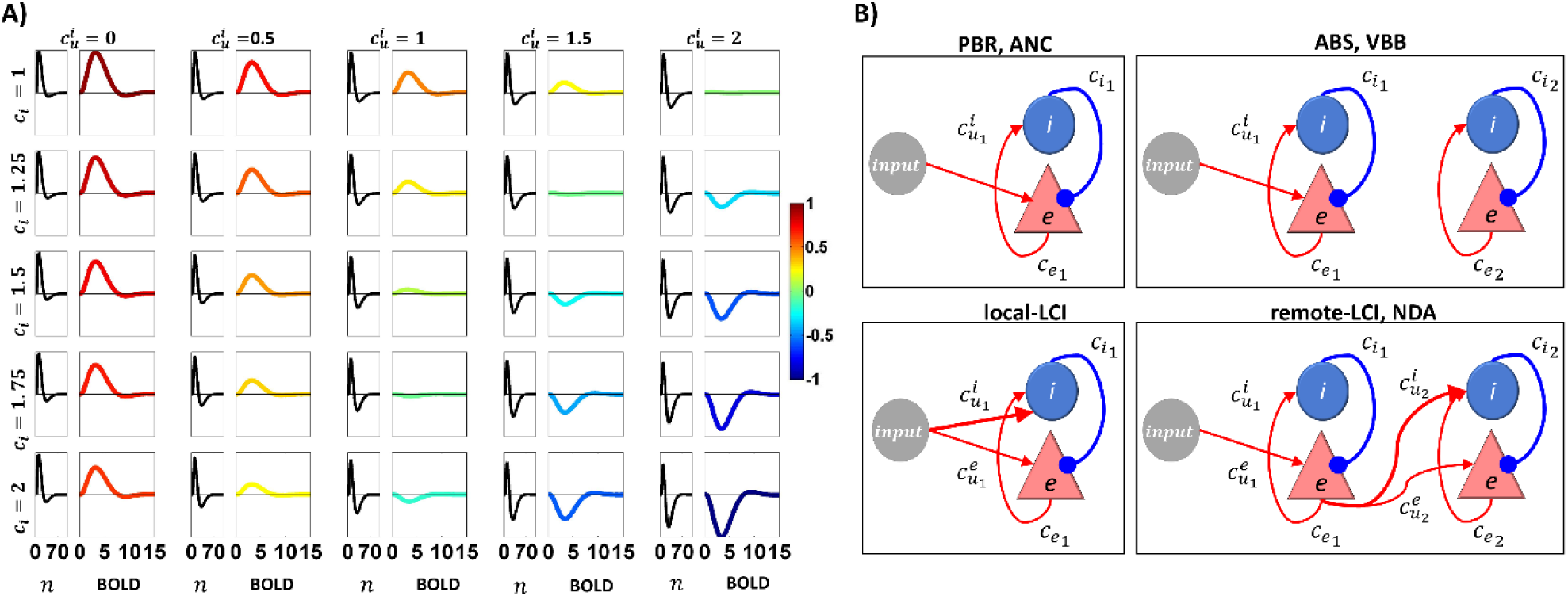
Figure 2. Inhibitory/excitatory balance for the different BOLD mechanisms. A) Effect of 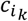 and 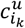, keeping 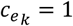 and 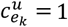, in a *k*–*th* region, on the total neuronal activity, i.e. 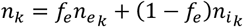 (first plot, black trace, unit: milliseconds), and the corresponding elicited BOLD response (second plot, colored trace, unit: seconds), after a very fast positive (delta-like) input. The amplitude of the BOLD responses was normalized to that of the top left corner, which corresponds to the typical situation in which a PBR is obtained from an excitation. The figure indicates that zero input to the inhibitory state yields no or small deactivation in the neuronal response following the activation and a consequent PBR. On the other extreme, a strong input to the inhibitory state yields a more bipolar neuronal activity profile and a consequent NBR, but only if the inhibitory to excitatory connectivity is stronger than the excitatory to inhibitory one. The strength of the inputs to both states must be unbalanced to generate BOLD responses comparable in amplitude to the typically modelled PBR. B) Configurations of connectivities in the neuronal model to simulate the mechanisms. Thin (thick) lines corresponds to the value 1 (2). These values are shown in Table 1. The self-connections, present in all models and states, were omitted for visual simplicity.

According to simulations, for the *k*^*th*^ region, the balance between 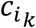 and 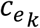, and between 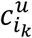 and 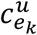 in equations (3) determines the polarity of the BOLD response. Figure 2 shows this polarity for the range 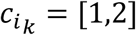 and 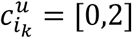, with 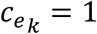 and 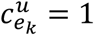, using the standard extended balloon model. Irrespective of the value of 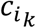, a PBR is obtained when 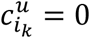; while an inhibitory-related NBR is obtained for 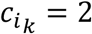 and 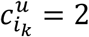. We use this value of 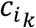 for all mechanisms since it is reasonable to assume that inhibitory gains are stronger than excitatory ones—inhibitory interneurons tend to have synapses closer to the soma (Megi´as et al., 2001; Villa and Nedivi, 2016). We use the PBR configuration in region 1, i.e. 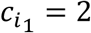 and 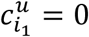, to simulate non-inhibition-related NBR mechanism, i.e. ANC, ABS and VBB. To simulate the local-LCI, we assume the input is acting on both states of region 1, i.e. 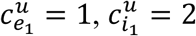, while this effect is not transmitted to region 2, i.e.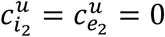. Obviously, the latter also holds for ANC, ABS and VBB. To simulate inhibition in region 2 for both remote-LCI and NDA, we set a configuration that generates positive neuronal response and PBR in region 1, i.e. 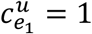 and 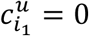, which is transmitted stronger to the neuronal inhibitory state of region 2, i.e. 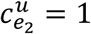 and 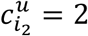. The configuration of neuronal connectivities for all mechanisms is summarized in Figure 2.

For the *k*^*th*^ region, except for NDA, times constants for the neuronal states are set to 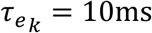, to obtain a fast-neuronal response (like an IED) when an ultra-short input pulse is applied, and 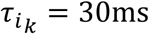 to simulate a slower inhibitory-related recovery of the neuronal activity (similar to the hyperpolarizing wave seen after IEDs). These time constants are in the range of those typically used in neural mass modelling literature involving excitatory and inhibitory populations of neurons (David et al., 2006; Robinson, 2005). For local-LCI, 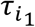 can be as high as 200ms to simulate the slower recovery wave reported after IEDs, c.f. (Pittau et al., 2013). We assume that, due to conflicting interactions between the different nodes of the affected network, the recovery of the neuronal activity in NDA is much slower than in other mechanisms, i.e. 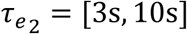.

For all mechanisms, we force the neuronal activity of both states to be higher than 40% (*n*_0_=0.4) of its baseline activity. Contribution of excitatory (inhibitory) activity to the generation of the BOLB response is set to 75% (25%), following the proportion of excitatory/inhibitory neurons in the human cortex.

Under normal conditions, the neurometabolic coupling gain can be set to *κ*=0.05 and the effect of the input on *g, c_B_* and *E* is little. For simplicity, we discard equations (14), (15) and (16) and set 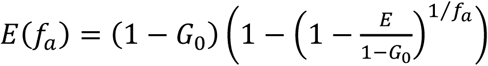 to simulate and estimate the model under LCI, NDA, ABS and VBB mechanisms.

For all mechanisms we disconnect the windkessels from the common supplying artery, i.e. *ρ*_*A*_=0, except ABS; and from the common draining vein, i.e. *ρ*_*V*_=0, for the VBB. This corresponds to vessels with very large diameters or regions far apart from each other, i.e. not sharing any vessel.

### Values for blood resistance and inertia in the artery and vein

We provide the rationale behind the proposed values for *ρ*_*a*_, {*ρ*_*X*_}*X*=*A*,*V* and *τ_X_* in

Table 2, where *X* denotes either artery or vein. Following (Mandeville et al., 1999) we propose that arteriole plus capillary resistance is around 65% of the total arteriole plus capillary plus windkessel resistance:

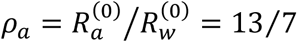

This value can be as low as 1.4, if we use the equivalent resistances used in VAN (Boas et al., 2008). However, we decide to account for any variability between the resistances by allowing *ρ*_*X*_ to vary. Its value is given by (see Appendix):

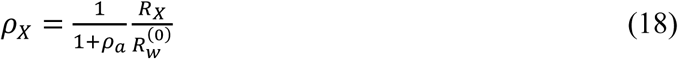

To calculate *R*_*X*_ we use Poiseuille’s formula:

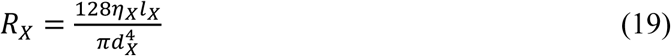

where *η_X_, d_X_* and *l*_*X*_ are the dynamic blood viscosity, the inner diameter and length of the vessel, respectively. For hematocrit of 45% and diameters of the final segments of the arteries (surface and penetrating arteries) ranging between 100—500µm (Bevan et al., 1999; Gutierrez et al., 2014), the viscosity of blood is *η_b_*∼2—3cP (Pries et al., 1992). If the length of these arterial segments is around *l*_*X*_∼1cm, *R*_*X*_ ranges between 0.3—917 mmHg/µL. Moreover, using the estimates in (Boas et al., 2008), the equivalent value of 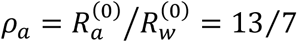 for the vascular network is ∼970 mmHg·s/*μL*. Since we are interested in rough estimates of the ranges, we shall use the same ranges for the initial segments of the veins. Evaluating these values in equation (18) yields the range *ρ*_*X*_∼0-0.33. We are interested in exploring ABS and VBB mechanisms with a significant NBRs. In Section “Predicted responses for single impulses and block designs” we show that for very small values of *ρ*_*X*_, the amplitude of the NBR is not significant. Thus, we restrict *ρ*_*X*_≥0.05 for the simulations and to establish the ranges allowed for the estimations in this paper.

To calculate the inertial time constant we use (Spronck et al., 2012):

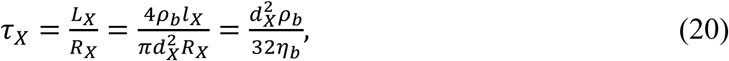

where *L*_*X*_ and *ρ*_*b*_ are the “blood inertia” (inertial force per unit of acceleration) and mass density, respectively. Using *ρ*_*b*_=1.05g/mL we obtain:

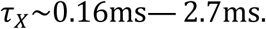

Our simulations (not shown in this paper) demonstrate that varying the value of this parameter within this range causes a negligible effect on the HRFs. Therefore, we fix the value to 1ms.

### Model simulations

Throughout the paper, we performed simulations of the temporal behavior of the state variables and the BOLD signal under realistic levels of noise (Chen and Tyler, 2008; Kasper et al., 2017) given an input function *u*(*t*). Consequently we used the local linearization (LL) scheme (Biscay et al., 1996; Carbonell et al., 2005), suitable for the integration of stochastic nonlinear state space models. We adapted the LL to account for the imposed lower bound of *n*_1,2_ and *g*_1,2_. It is important to note that our state equations constitute a system of stochastic ordinary differential equations (SODE) with a singular mass matrix **M** (see Appendix). Unfortunately, the LL integration scheme cannot cope with this type of SODE systems. To circumvent this problem, we “regularize” the mass function matrix with the substitution 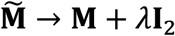, with a value of *λ* much smaller than the least singular value of **M**. We then pre-multiply the drift term by 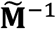 to eliminate the mass function matrix. To evaluate the effect of this modification we simulated the unnoisy modified ODE system using LL and compared it with the original ODE with the mass matrix function using MATLAB R2013b’s *ode15s* (Shampine and Reichelt, 1997). We could verify that the results were almost identical (comparison not shown here). We therefore used the modified SODEs for the simulations.

Throughout the paper, the external input *u*(*t*) is modeled as a single or a train of very short pulses. Such pulse can be viewed as a fast depolarization of the neurons, a fast drop in their firing threshold, etc:

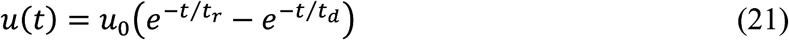

where *t*_*r*_=5 ms and *t*_*d*_=2.5 ms are the rise and decay times of the pulse, respectively, and *u*_0_ is adjusted to provoke a maximum amplitude of 1% BOLD signal change when generating a typical PBR (Friston, 2002; Logothetis et al., 2001; Zheng et al., 2002) (see Figure 2 and

Table 2). In order to avoid the system to blow up (and provoke an unrealistic change in neuronal response) when using a train of *P* pulses (especially for a very low inter-stimulus-interval (ISI) or high pulse frequency), we choose:

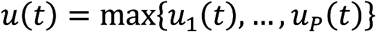

### Detection of mechanisms using the general linear model

The fastest and most widely-used method to detect significant BOLD responses is the GLM regression (Friston et al., 1995):

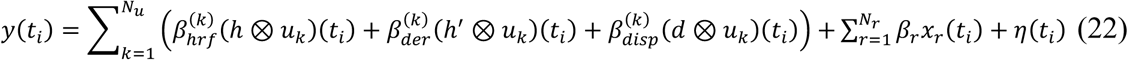

where {*t*_*i*_}_*i*_=**,…,N**, being *N* is the number of scans, *N*_*u*_ is the number of types of inputs 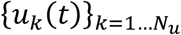,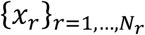 are the confounding or nuisance regressors (e.g. the motion parameters) and *β_r_* are their effect sizes; *h, h*^′^ and *d* are the canonical BOLD HRF (Glover, 1999), its temporal derivative and its dispersion, respectively. The noise is pre-whitened by applying a first order autoregressive (AR) model, i.e. (AR(1)), to the signal.

To demonstrate that the NBR mechanisms are detectable using GLM, we simulated a set of 10-minute BOLD signals with a single type of input (*N*_*u*_=1) consisting of a random Poisson train of pulses with average frequency of 2.6/min. This input was the same across all models and trials. We then created a simulated set of *N*=300 fMRI scans with repetition time *T*_*R*_=2s by adding the simulated BOLD signals to the voxels of a real EPI image. The trials from the same type of model were added to neighboring voxels to form spatial clusters. The amplitude of the simulated BOLD signal in each cluster was multiplied by a Gaussian spatial kernel with the maximum at the center of the cluster. The resultant set of images was further corrupted with colored noise, according to a spectral density given by 1/*f*^*p*^, with 0 < *p* < 1, to account for biological noise inherent to the BOLD signal but not attributable to the temporal filtering of the hemodynamic response function (HRF) (Chen and Tyler, 2008). No nuisance regressor was included in these simulations, i.e. *x*_*r*_ = 0. Following standard fMRI preprocessing procedures, the simulated fMRI scans were spatially smoothed with a Gaussian kernel of 8 mm FWHM. For each voxel, the vector of coefficients ***β*** = [*β_can_ β_der_ β_disp_*]*^T^* of the GLM in (22) were estimated using SPM (Friston et al., 1995). Using an F-contrast, we selected the voxels where the null hypothesis, **I**_3_***β***=**0** was rejected with p<0.05, after correcting for multiple comparisons using the Family-wise error criterion, i.e. the significant voxels.

### Estimation of Hemodynamic Response Functions

Although the responses fitted by (22), i.e. the first order Volterra kernel (Friston, 2002), can reconstruct a large variety of HRFs, we used a more adaptable method to accommodate all possible extreme HRF waveforms, the near-neighborhood AR with exogenous variable method (NN-ARx) (J. Riera et al., 2004), which has been used to extract the HRF from fMRI time series. It is suitable to deal with the colored nature of the fMRI time series and the spatial correlation between neighboring voxels in the images. This linear method also allows for the incorporation of different types of inputs. By minimizing the Akaike information criterion, NN-ARx estimates the orders and coefficients of the AR, the order of a polynomial modeling drifts, and the delay of the HRF onset. With these, the HRF can be constructed (J. Riera et al., 2004). By applying the NN-ARx to the unsmoothed fMRI scans, we extracted the HRF in the significant PBR and NBR voxels of the GLM test.

### Classification of mechanisms

Once the significant voxels are detected and their HRF are estimated, it is necessary to classify them according to the type of NBR mechanism. For this purpose, we propose to build a classifier based on a machine learning algorithm. We simulated *M*=50 trials of 10-minute fMRI signals with *T*_*R*_=2s with random inputs consisting of trains of short pulses Poisson-distributed in time. To account for the entire span of HRF waveforms, we model the possible intra-individual and inter-individual variability in the parameters by randomly sampling their values, for each trial, from a uniform distribution within the intervals specified in

Table 2. With the NN-ARx-based HRFs for all trials, normalized by the maximum of their absolute value, we formed a 6*M*×*N*_*T*_ matrix of features. The rows number equals the number of trials times the number of classes, i.e. PBR, NDA, ANC, ABS, VBB and LCI. The number of columns is the number timepoints of the HRF, i.e. *T*/*T*_R_=32/2=16. This matrix of features and the corresponding vector of classes were used to create a machine learning ensemble classifier (McLachlan, 2004; Seber, 1984). The classifier was based on quadratic discriminant analysis to account for distributions of the features with different means and variances among classes.

### Validity of the use of linear regression methods

BOLD responses are non-linear with respect to the input (Buxton et al., 2004; Friston et al., 2000). Thus, GLM and NN-ARx—both linear regression methods—have to be used with caution. Under the assumption of the extended balloon model, the validity of the GLM was investigated by quantifying the effect size of second order Volterra kernels (Friston et al., 2000). Under the same model, the validity of the NN-ARx to detect and estimate HRFs was evaluated in (J. Riera et al., 2004). In general, for balloon/windkessel models—with typical canonical-like responses (Glover, 1999)—these linear models are suitable for spatial detection estimation of the HRF. However, in this paper we introduce important modifications of the balloon/windkessel model, in particular related to significantly nonlinear flooring effects due the lower bounds imposed to the states variables *n*_±_ and *g* and long recovery times of the BOLD response.

To explore the effect of these nonlinearities, we compared the simulated BOLD response of a train of periodic short pulses (the nonlinear response), say *y*_*train*_(*t*), with the sum of the BOLD responses for the same pulses, presented individually (the linear response), say *Y*_*sum*_. We calculated two measures: a) the temporal correlation between *Y*_*train*_(*t*) and *y*_*sum*_(*t*), and b) given the areas under the curves 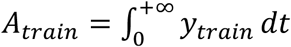 and 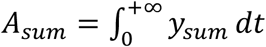, their relative absolute difference 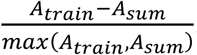. This was done for pulse train frequencies of 0.03, 0.05, 0.1, 0.2, 0.3, 0.5, 1, 1.5, 2 and 3 Hz, to determine for what ISI the nonlinearities become significant. This establishes a criterion for the assumption of linearity and furnishes the ability to propose correction methods when such nonlinearities are present.

### Estimation of the parameters of the models

Most of the parameters of the models have been estimated previously in both simulations and real data. For example the parameters *ε, *τ*_s_, *τ*_f_, *τ*_MTT_, β, E*_0_ were estimated in (Friston, 2002; J. J. Riera et al., 2004); while *τ_w_* and *b* were estimated in (Zheng and Mayhew, 2009). In addition, both the neurovascular and neurometabolic coupling gains, *ε* and *κ* were estimated from CBF data in normal and epileptic rats in (Song et al., 2016). We investigated the accuracy and precision in the estimation of the rest of the newly introduced parameters, determinant in the formation of the NBRs. These are *τ_i_*, 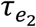, *κ*, *ρ*_*A*_ and *ρ*_*V*_. To that end, we performed simulations and estimated their values restricted to the ranges provided in

Table 2. For each mechanism, we simulated 3500 random realizations of a 60-second BOLD signal. The input for each trial was a random set of unit short pulses distributed across the interval [5s 40s] following a Poisson distribution. The simulated BOLD signal was sampled at *T*_*R*_=2s. For parameter estimation, we applied a local linearization filtering for nonlinear state space models (Jimenez and Ozaki, 2003) and calculated the likelihood distribution of the innovations. Using MATLAB R2013b’s *fmincon*, we estimated the values of the parameters, alongside the standard deviation of the observation noise parameter ζ, that maximized this distribution. This method was also used to estimate the parameters in the real data.

### EEG-fMRI data

The experiments were carried out at the Nicklaus Children Hospital, Miami, FL, in accordance with the World Medical Association declaration of Helsinki—ethical principles for medical research involving human subjects. The protocol of the study was approved by an Institutional Review Board (WIRB # 20160218). Parents or legal guardians of patients signed a written approved-informed consent.

We acquired four 10-minute trials of simultaneous EEG-fMRI data from a set of pediatric patients (age 12-17) with focal epilepsy. Using the fMRI trials for which the IEDs were better identified from the EEG we fitted GLM and NN-ARx to detect and calculate IED-related PBRs and NBRs. The IEDs were visually detected and classified into several subtypes by two experts based on their morphology, the semiology of the patient and other neuroimaging modalities (e.g. stereological EEG, PET and SPECT, when present). The different subtypes of IEDs and other types of events were used as different types of inputs (conditions) in GLM and NN-ARx. Additionally, motion correction parameters were included as nuisance regressors. We then attempted to classify them into NBR mechanisms using the machine learning classifier. Finally, we estimated the parameters of the corresponding particular sub-model.

Using a Philips 1.5T scanner with a 16 channel SENSE Rx coil, fMRI was acquired using a GE-EPI sequence. Each scan consists of 21 interleaved slices 6 mm-thick with 2 mm gap, in-plane voxel size of 3×3 mm and FOV = 204 mm. Flip angle (FA) was 90°, repetition time *T*_*R*_=2 s and echo time *T*_*E*_=45 ms. For the purpose of anatomical reference, a high resolution T1-weighted image was acquired using a spoiled 3D-GRE with *T*_*R*_ = 9.7 ms, *T*_*E*_ = 4 ms and FA = 12°. The structural MRIs have 90-100 slices, covering the whole brain. In some cases, a T2-weighted 3D image was also acquired with parameters: *T*_*R*_ = 25 ms and *T*_*E*_ = 3.732 ms, NAX=1, FA = 30°, FOV = 240 mm and 160 2 mm-thick axial slices. The fMRI volumes were preprocessed using SPM (http://www.fil.ion.ucl.ac.uk/spm/). They were corrected for motion correction and spatially smoothed with a 8-mm Gaussian kernel. Both smoothed and unsmoothed images were used in GLM analysis to detect significant voxels. Although the minimum variance estimator of GLM may be biased due to non-Gaussian noise (Friston, 2007), the latter was necessary to detect near PBR and NBR, as it was suggested by (Goense et al., 2012; Harel et al., 2002) for detecting the NBR of vascular origin. The GLM analysis and the results were masked to the gray matter using the SPM segmentation obtained from the anatomical image (Ashburner and Friston, 2005).

In order to detect the IEDs, EEG was recorded using a 10-10 system 32-channel EasyCap (BrainAmp MR, Brain Product GmbH) simultaneously with the fMRI. We used MRI-compatible EEG amplifiers (BrainAmp MR, Brain Product GmbH). EEG signals were sampled at 5 kHz and digitized (0.5μV resolution) (BrainVision Recorder 1.4, Brain Products GmbH). The majority of EEG electrodes had impedances lower than 5 kΩ. The electrocardiogram (ECG) was measured with an ECG electrode attached to the middle of the back of the patients. To synchronize the EEG with fMRI scans a trigger marking the beginning of the scans was sent to the EEG recording laptop. To ensure the highest temporal precision, the clock of the laptop was synchronized to the 10MHz clock of the MR-console using a Syncbox (Brain Products GmbH). The following EEG preprocessing was performed using BrainVision Analyzer 2 (Brain Products GmbH). To remove the MR-related artifact, the EEG data was first subsampled to 50 kHz using *sinc* interpolation to virtually increase its resolution and correct the random phase jittering—of no less than 0.2ms resolution determined by the 5kHz sampling rate—that is present in the scan markers. This phase jittering has significant negative effects in the estimation of the MR gradient artifacts since the latter can change as fast as 0.2ms. Subsequently, we applied a method for removing the MRI gradient artifacts (Allen et al., 2000), based on the estimation of an average artifact template. The resultant EEG data was bandpass filtered within 0.5-125 Hz. After marking the R-waves using a semiautomatic tool, we applied a method to detect and remove the effects of the balistocardiogram (Allen et al., 1998). Finally, we applied ICA based on the infomax method (Bell and Sejnowski, 1995; Makeig et al., 1997) to remove further artifacts.

Since IEDs last about 70-200ms, the input *u*(*t*) is modeled as a train of short pulses. To account for the actual time in which the slice containing the region with significant voxels was acquired, the inputs *u*(*t*) were transformed according to *u*(*t+nT_R_N_Z_/*), where *n* is the position of the slice according to the sequence in which they were acquired and *N*_*Z*_ the number of slices the whole scan.

## Results

### Predicted responses for single impulses and block designs

We explored the existence and behavior of NBR mechanisms under two types of inputs: single short pulses, e.g. event-related stimuli or IEDs, and sustained/periodic simulation, e.g. block-design stimulation paradigms. Figure 3 shows the predicted responses for the mechanisms proposed in this paper, and their sensibility to the relevant parameters, following a short single event. Note that the NBR is not significantly affected by the values of *τ*_*i*_. Thus, it cannot be solely estimated from BOLD signals and combined EEG-fMRI-based estimations are needed. This was already demonstrated in (Riera et al., 2007). Figure 4 shows the responses for a sustained input. Contrary to the block-design, the nonlinear flooring effect of the neuronal activity is not expected to happen in the NBR region following a single short event. As expected, the recovery of the network is reflected in the NBR. Regarding the vascular phenomena, the NBR only occurs in the presence of blood resistance in the shared vessel. The higher the resistance, the higher the amplitude of the NBR. In fact, simulations not shown demonstrate that for negligible blood resistance the amplitude of the NBR is also negligible in comparison to the PBR. For ANC, a disproportionately high neurometabolic to neurovascular coupling ratio yields NBR. This is not sustained for continuous stimulation (or very high frequency pulse stimulation) during several seconds—consequence of the nonlinear flooring effect of the tissue O_2_ concentration. If stimulus frequency in block-designs is decreased, this nonlinear effect become less determinant and the NBR can occur (see Figure 5).

Table 2. The light blue arrow indicates how the NBRs change by increasing the value of the parameters. Besides BOLD responses, we also show other candidate observables in MRI: rCBF and rCBV. Expectedly, the higher the recovery time constant in the NDA mechanisms, the slower the recovery of the NBR. We also note that, higher neuro-metabolic coupling gain yields higher NBR amplitudes in the ANC mechanism. Besides, the higher the arterial and vein resistance, relative to the arteriole and venule compartments, in the ABS and VBB mechanisms, respectively, the higher NBR amplitude.

**Figure 3.**
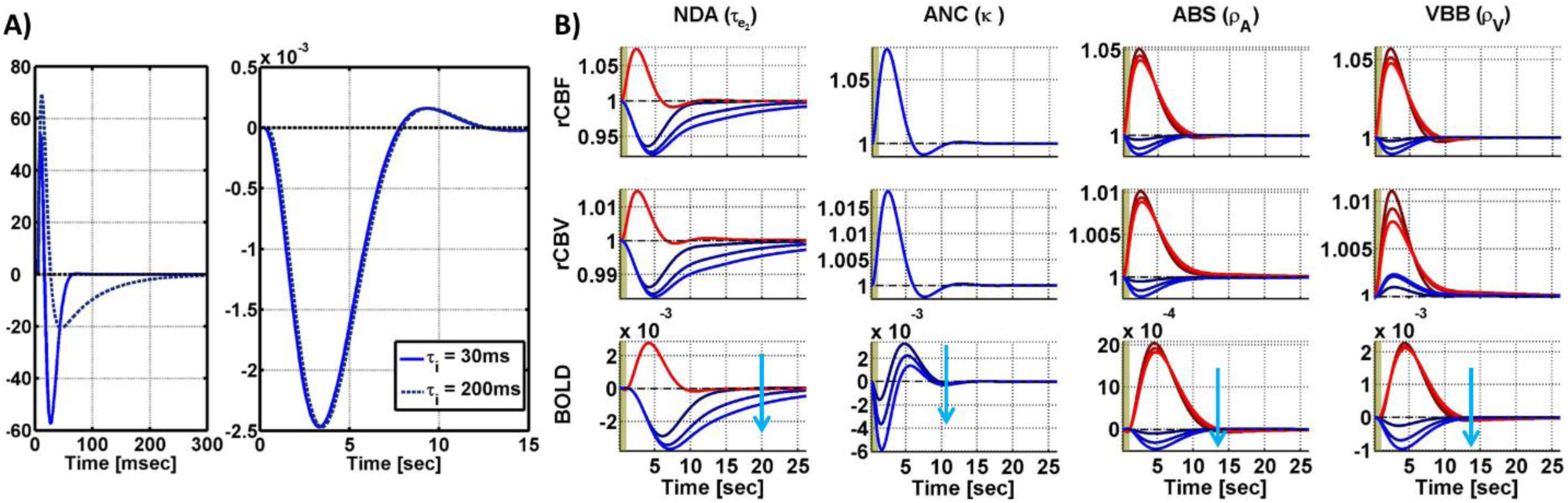
Simulation of the LCI, NDA, ANC, ABS, and VBB mechanisms after a single short pulse. Red(blue) corresponds to PBR(NBR). A) Neuronal response (in percentage, first plot) to a short pulse in LCI and consequent BOLD response (second plot) in LCI for the extreme values of *τ*_*i*_. The effect *τ*_*i*_ on the NBR in LCI is negligible. B) Each column corresponds to a parameter that significantly determines the NBR waveform. The sensibility of the responses to these parameters is illustrated with different curves corresponding to three different values of the parameters covering the ranges in

**Figure 4.**
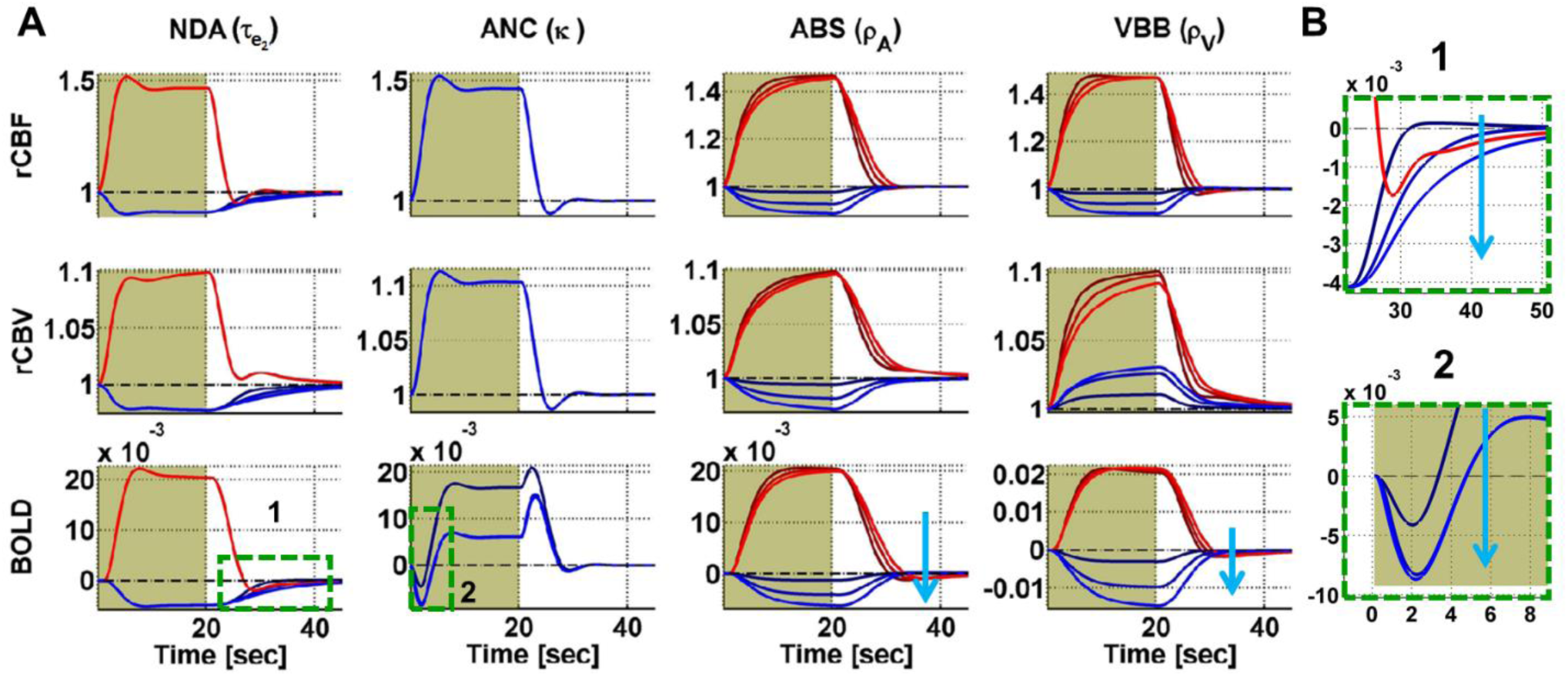
A) Simulation of the NDA, ANC, ABS and VBB mechanisms after a 20s block stimulus. For better visualization, some parts of the responses (labeled as 1 and 2) are magnified and shown in the insets in B). Note that, for a block design, there is a lower bound for the NBR for the NDA mechanism due to the flooring effect of the neuronal activity. The sensitivity of the NDA NBR recovery shape to the recovery time constant is less noticeable for block designs than for single short pulses. Besides, for a block design, there is no sustained NBR for the ANC mechanism due to the flooring effect in the tissue O_2_ concentration, which cannot be less than zero.

**Figure 5.**
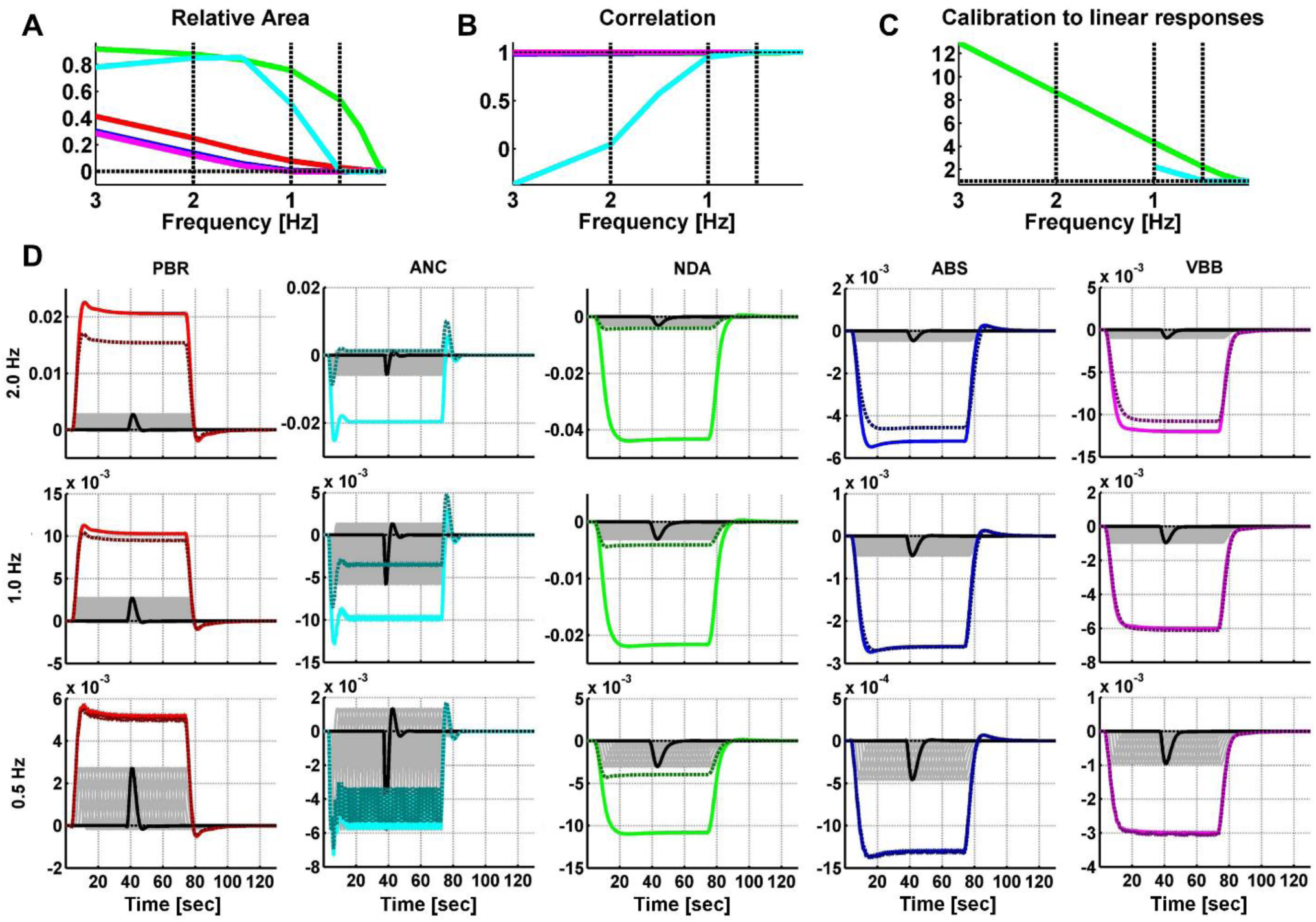
Empirical validity and correction of linear methods for detecting and estimating the HRF. A) Relative difference of the area under the curve between the nonlinear simulated response using the full periodic input and the linear addition of the simulated responses using each individual pulse separately, as a function of pulses frequency. For PBRs, the nonlinear effects become apparent for frequencies higher than 1Hz (below 1s of pulse separation); for NBRs in ABS and VBB above 1.5 Hz, and NBRs in ANC, above 0.5 Hz. For NDA the response is nonlinear for all frequencies. B) Correlation between both responses. We note that all nonlinear PBRs and NBRs are practically scaled versions of their corresponding linear compositions, except for the NBR in ANC above 1 Hz. Since they only differ by a sign, the results for PBR also apply for LCI. C) We calculated “scale calibration curves” to account for scale differences between the nonlinear and linear responses. These curves were used to calculate a scaling time series, as a function of the average time-dependent frequency profile of the inputs *u(t)*, to correct the nonlinear flooring effects in NDA and ANC. Application of this methodology to ABS or VBB signals was deemed unnecessary since the frequency on inputs, whether in event-related or block-design experiments, is usually below 2 Hz, as it is the case of the IEDs in our real data. D) Illustration of the effect of the nonlinearities in PBR and NBR for three values of pulse frequency. The dark(light) color trace represents the nonlinear(linear) response. The individual pulses are represented with the gray curves. For visual clarity, we highlight one of these individual responses in black. With the purpose of clear visual identification, we associate each mechanism with a distinct color throughout the paper: PBR (red), NDA (green), ANC (cyan), ABS (blue) and VBB (magenta).

### Accounting for nonlinear effects

Figure 5 illustrates how the effect of the nonlinearities on the responses was assessed and how it was accounted for in the linear methods used in this paper. We evaluated the relative absolute value of the difference between the nonlinear and linear simulated responses defined in “Validity of the use of linear regression methods” as a function of frequency. For these simulations the values of parameters were set to the average of the ranges in

Table 2. Note that the response becomes notably nonlinear for ISI less than 1.5s for ABS-NBR and VBB-NBR; less than 1s for PBR and less than 2s for ANC-NBR. For the frequencies analyzed, ISI is lower than or comparable to the recovery time constants in NDA; thus, its responses are always nonlinear. To investigate if the differences in areas were due to either scale or shape differences between the responses, we calculated their correlation. Except the ANC-NBR 1Hz (consistent with the block-design simulation in Figure 4), all pairs only differed by a scale factor. This means that we can use the linear regressions methods if we properly multiply the fMRI time series at each time point by a correction scale which would depend on the “instantaneous” frequency of the input. We calculated the temporal frequency profile of the input using a moving average window. Using this temporal frequency profile and the calibration curve we calculated the scaling time series that was multiplied by the fMRI time series to correct the nonlinear effects. Since neither the frequency of the simulations in this paper, nor the IEDs detected in the real data have a frequency higher than 1 Hz, we use this approach only for the analysis of ANC and NDA mechanisms. The “calibration curve” specifying the value the fMRI signal has to be multiplied, as a function of frequency, is also shown in Figure 5.

### Detection, estimation and classification of NBR mechanisms

Figure 6. Detection, using GLM, of simulated spatial distribution of PBRs and NBRs in a realistic sequence of echo-planar fMRI scans. A) 3D glass brain showing the regions where the mechanisms where simulated. B) F contrast (the 3-order identity matrix) and design matrix of the GLM, used to detect significant voxels. C) 2D glass brain showing the F-statistics. The plots show the responses predicted by the GLM (color curves) and the adjusted data (black dots), for each mechanism. Beside each plot, the estimated values and confidence intervals of the coefficients, *β_can_, β_der_* and *β_disp_*, of the GLM are shown. PBR (red), NDA (green), ANC (cyan), ABS (blue) and VBB (magenta).

**Figure 6.**
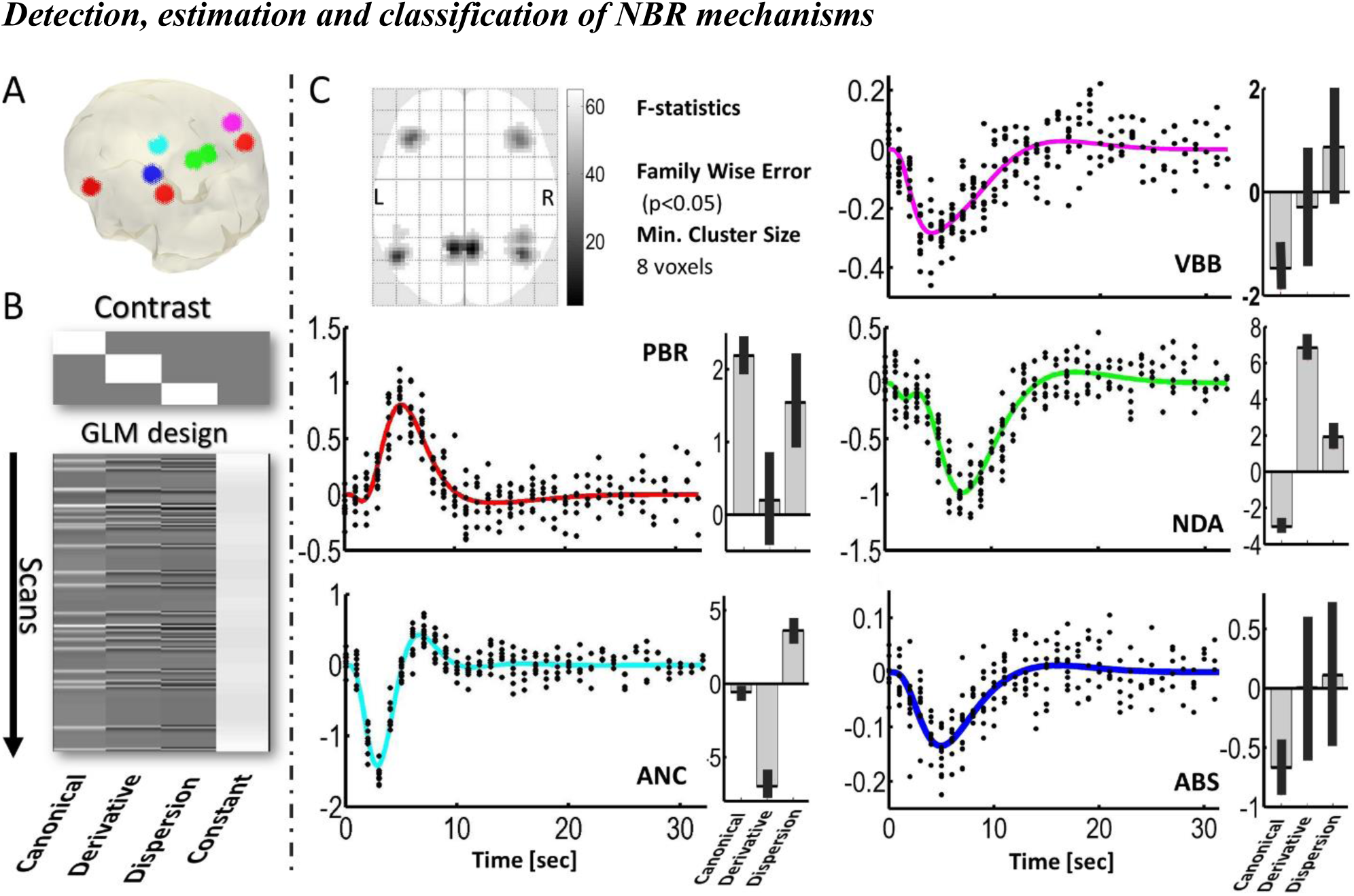
Detection, using GLM, of simulated spatial distribution of PBRs and NBRs in a realistic sequence of echo-planar fMRI scans. A) 3D glass brain showing the regions where the mechanisms where simulated. B) F contrast (the 3-order identity matrix) and design matrix of the GLM, used to detect significant voxels. C) 2D glass brain showing the F-statistics. The plots show the responses predicted by the GLM (color curves) and the adjusted data (black dots), for each mechanism. Beside each plot, the estimated values and confidence intervals of the coefficients, *β_can_, β_der_* and *β_disp_*, of the GLM are shown. PBR (red), NDA (green), ANC (cyan), ABS (blue) and VBB (magenta).

Figure 6 indicates that it is possible to detect voxels which are significantly exhibiting simulated NBR mechanisms using GLM. The mechanism with the least significance is ABS—because its NBR has the least amplitude.

Figure 7 illustrates how we built the machine learning classifier of mechanisms based on the quadratic discriminant analysis of ensembles of NN-ARx HRFs—estimated from simulated fMRI signals with random values of the parameters accounting for intra-individual and inter-individual variability, given ranges in

**Figure 7.**
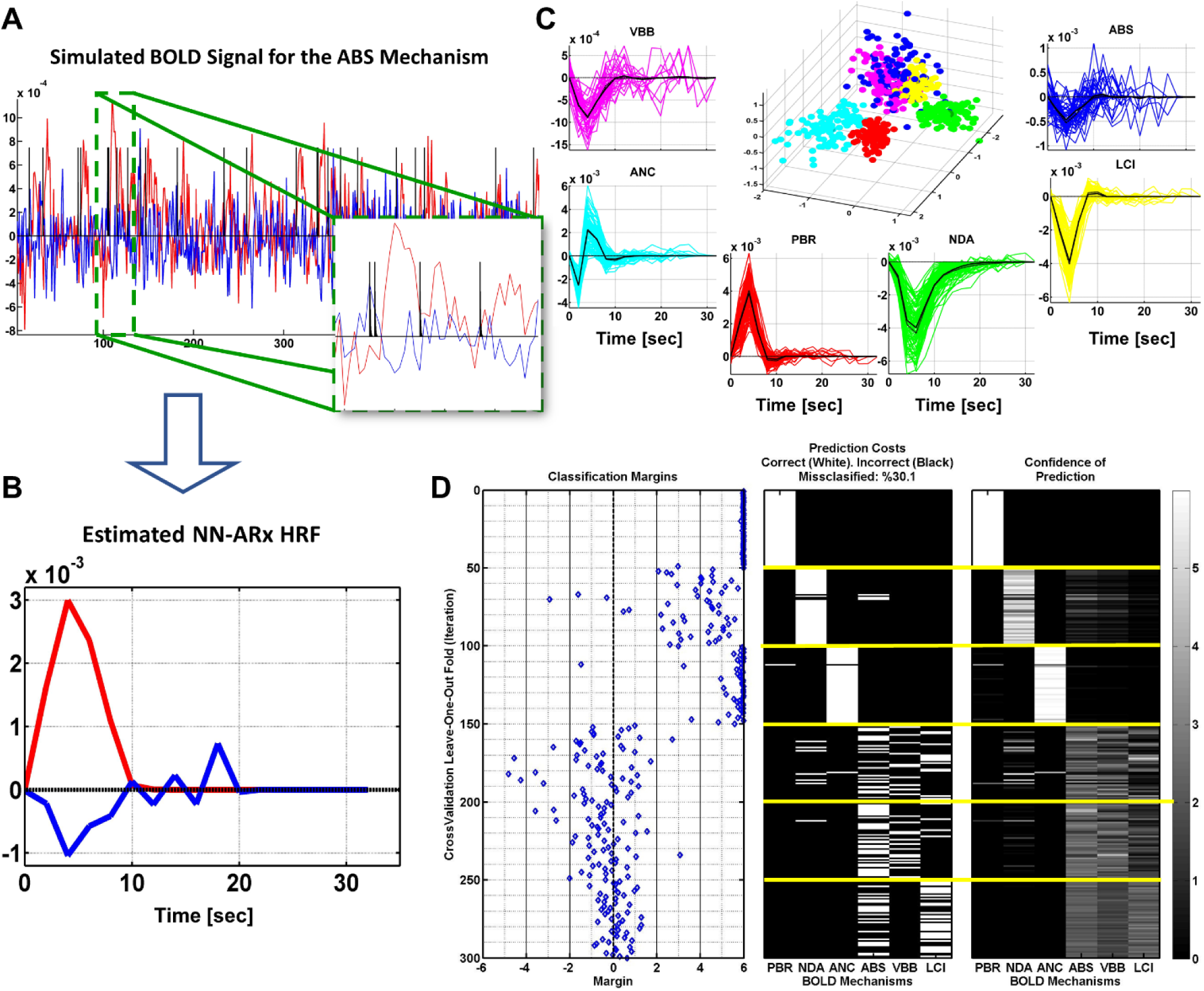
NN-ARx estimation and classification of HRFs. A) Example of a trial of simulated fMRI time series for the ABS mechanism. The input *u(t)* is represented with a black trace. B) HRFs estimated using NN-ARx from the example in A). C) Estimated HRFs from all trials. The black continuous curve represents the average across trials; while the black dash curve corresponds to a simulation without noise. To geometrically illustrate their separability, the 3D plot depicts the scores of the first 3 components of the PCA decomposition of the matrix formed by stacking all HRF as row vectors, after normalizing their amplitudes. PBR (red), LCI (yellow), NDA (green), ANC (cyan), ABS (blue) and VBB (magenta). D) Results of the one-fold cross-validation of the machine learning classifier. The first plot shows classification margins, i.e. the difference between the classification confidence for the true mechanism and the maximal classification confidence for the false mechanism. The second plot shows the prediction costs. The third plot shows the confidence in the prediction. As expected, the PBR is distinguishable from the NBRs. NDA and ANC mechanisms can be classified in general; while LCI, ABS and VBB might not.

Table 2. We also demonstrate the ability of this classifier to predict new mechanisms using one-fold cross-validation, i.e. leaving one HRF out for prediction and using the rest as the training set. Unsurprisingly, the PBR response, which we included as reference, is clearly separable from the NBRs. The ANC is distinguishable from the PBRs, even though its HRF can have a significant positive overshoot for 0.15<κ<0.3. NDA and ANC can be distinguished from each other and from the rest of the mechanisms, owing to the prolonged recovery of the former and the fast and bipolar shape of the latter. However, the margin of classification and the confidence of prediction of LCI, ABS and VBB are very low. This means that they cannot be distinguished from each other, at least merely from fMRI signals. Mechanisms were incorrectly classified in around 30% of the cases, mostly among LCI, ABS, VBB and some NDA with low 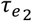.

Figure 7. NN-ARx estimation and classification of HRFs. A) Example of a trial of simulated fMRI time series for the ABS mechanism. The input *u*(*t*) is represented with a black trace. B) HRFs estimated using NN-ARx from the example in A). C) Estimated HRFs from all trials. The black continuous curve represents the average across trials; while the black dash curve corresponds to a simulation without noise. To geometrically illustrate their separability, the 3D plot depicts the scores of the first 3 components of the PCA decomposition of the matrix formed by stacking all HRF as row vectors, after normalizing their amplitudes. PBR (red), LCI (yellow), NDA (green), ANC (cyan), ABS (blue) and VBB (magenta). D) Results of the one-fold cross-validation of the machine learning classifier. The first plot shows classification margins, i.e. the difference between the classification confidence for the true mechanism and the maximal classification confidence for the false mechanism. The second plot shows the prediction costs. The third plot shows the confidence in the prediction. As expected, the PBR is distinguishable from the NBRs. NDA and ANC mechanisms can be classified in general; while LCI, ABS and VBB might not.

### Accuracy and precision of the model parameters

Figure 8 depicts the results of the estimation of the most relevant parameters in this paper, i.e. those determining the amplitude and shape of the NBRs. These are *τ*_*e*2_, *ρ*_*A*_, *ρ_A_*, *ρ*_*V*_ and *κ* for NDA, ABS, VBB and ANC, respectively. The figure contains a plot for each mechanism in which each point is a pair of estimated vs simulated values of the parameters. Each plot contains 3500 points corresponding to fMRI 60-second trials with random inputs and random values of the parameters within the ranges in

**Figure 8.**
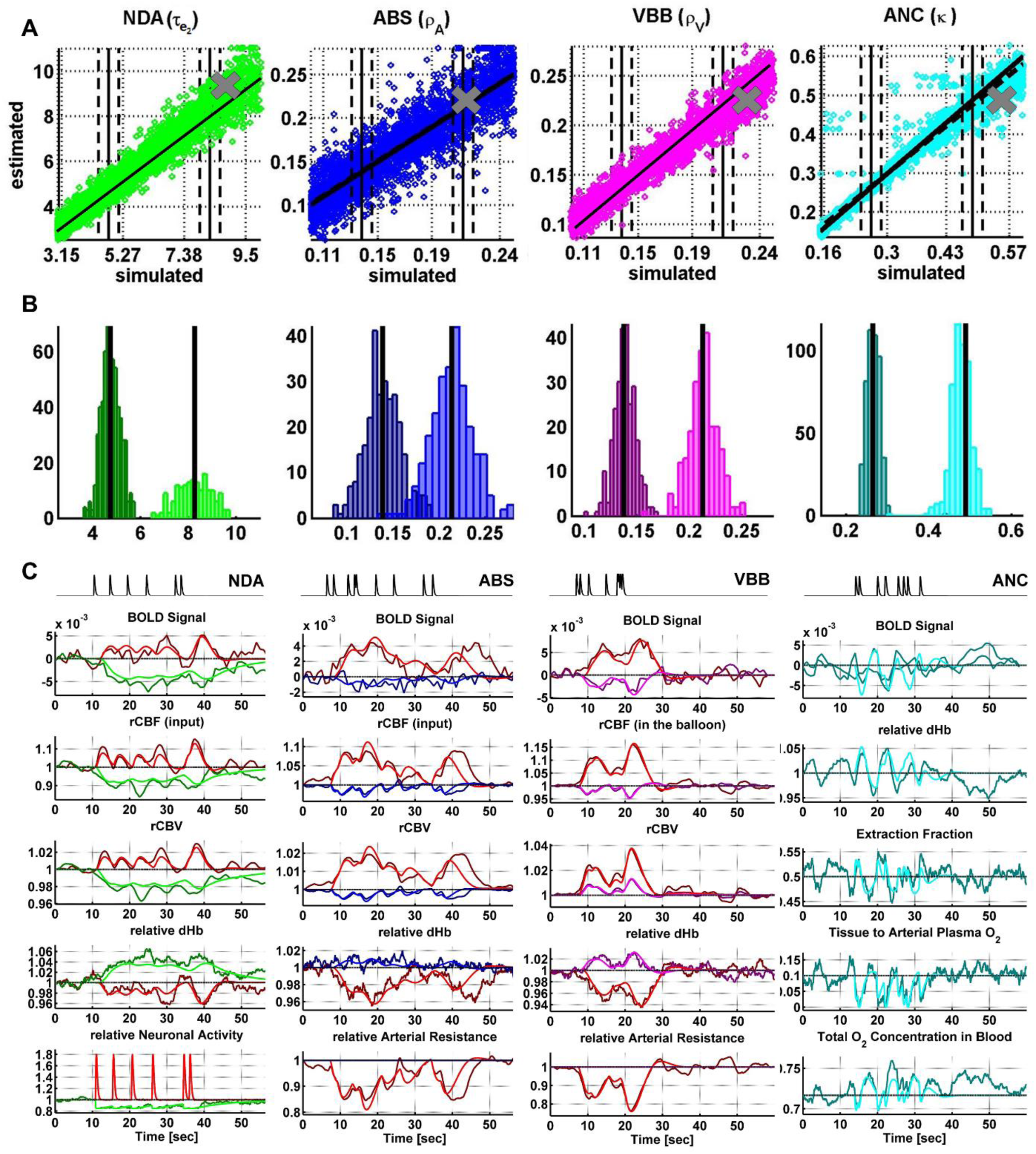
Results of the estimation of the parameters that determine the NBRs for NDA, ABS, VBB and ANC mechanisms. A) Each plot shows 3500 points, each corresponding to the pair (simulated, estimated) value of the parameter. The black lines represent the slopes of linear regressions. Neither the slopes were significantly different from 1 nor the intercept from 0 (*p∼*0). Thus, since the distribution of estimated parameters is approximately Gaussian (see B)), the estimation is accurate and unbiased. B) To illustrate the precision of the estimation, we show the histograms of the estimated values within the dash band in the corresponding plot in A). C) For the points represented as thick crosses in each plot in A), we also illustrate the temporal behavior of the BOLD signal and the most relevant state variables signal of the models used to simulate the mechanisms. PBR(red), NDA (green), ANC (cyan) ABS (blue) and VBB (magenta).

Table 2. To assess the performance of the estimation, we estimated the slope and the intercept of the line fitting the points, which were not significantly different from 1 and 0, respectively. Besides, Kolmogorov-Smirnov tests indicated the distributions of the estimated values were not significantly different from Gaussians with means coinciding with the simulated value. This means that the expected estimated value is equal to the simulated parameter. In other words, it is possible to calculate accurate (unbiased) estimates of the parameters. We evaluated the precision in the estimation of a parameter, i.e. the minimum significant difference between two estimated values, as 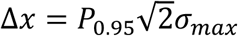, where *P*_0.95_=1.6449 is the one-tail Z-score for *p*<0.05 and σ_*max*_ is the maximum variance of the parameter within its range of estimation. The values of σ_*max*_ were 1.04s, 0.18, 0.14 and 0.24s^−1^ and the precisions were 1.9s, 0.06, 0.03 and 0.09s^−1^ for *τ*_r__, *ρ_A_, *ρ*_v_* and *κ*, respectively.

Figure 8. Results of the estimation of the parameters that determine the NBRs for NDA, ABS, VBB and ANC mechanisms. A) Each plot shows 3500 points, each corresponding to the pair (simulated, estimated) value of the parameter. The black lines represent the slopes of linear regressions. Neither the slopes were significantly different from 1 nor the intercept from 0 (*p*∼0). Thus, since the distribution of estimated parameters is approximately Gaussian (see B)), the estimation is accurate and unbiased. B) To illustrate the precision of the estimation, we show the histograms of the estimated values within the dash band in the corresponding plot in A). C) For the points represented as thick crosses in each plot in A), we also illustrate the temporal behavior of the BOLD signal and the most relevant state variables signal of the models used to simulate the mechanisms. PBR(red), NDA (green), ANC (cyan) ABS (blue) and VBB (magenta).

### Examples of possible NBR mechanisms in real data

In this section we present the results found in three cases of the dataset described in “EEG-fMRI data”. For each case we applied the following pipeline: a) GLM-based detection of voxels significantly correlated with the IEDs; b) selection of the region-of-interest (ROI) with high statistical significance; c) estimation of the NN-ARx HRF, averaged inside spheres within the ROIs; d) classification of the mechanism using the machine learning classifier—but complemented with the other criteria, such as the anatomical location of the clusters and the local gray matter properties; and e) estimation of the parameters of the sub-model associated with the identified mechanism. To estimate confidence intervals for the HRF, we estimated the empirical distribution of the null hypothesis of no significant response using a permutation test. This was done by estimating the NN-ARx HRFs from 5000 trials with random order of the IEDs. For each time point, the lower and upper confidence values were the 5 and 95 percentiles of this null HRF distributions, respectively.

The first example is unarguably NDA of Resting State Networks (RSN)—mainly the Default Mode Network (DMN)—during IEDs. All nodes of the DMN, as well as the Superior Frontal Gyri (SFG), the Middle Frontal Gyri (MFG), the left Inferior Frontal Gyrus (IFG), part of the right IFG, the right Fronto-Opercular region and the right Caudate Nucleus, were highly significantly deactivated. In addition, PBR was detected in the right IFG, which could be one of the foci of the IEDs, based on the semiology of the patient and their proximity to the EEG electrodes used to detect the IEDs—76 spikes were seen with highest amplitude in electrode F8. Other types of events were also marked and used as regressors in the linear models.

**Figure 9.**
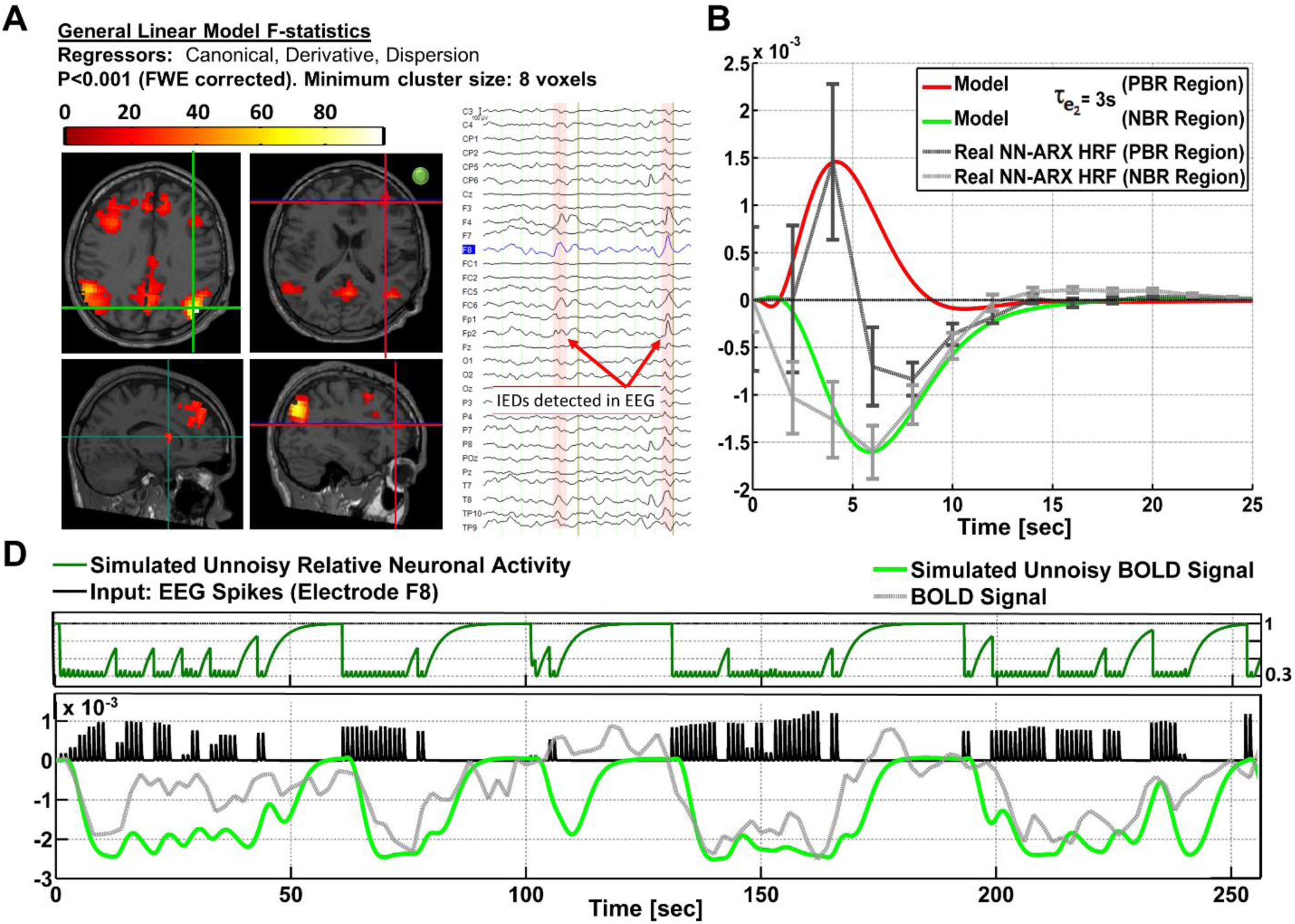

Figure 9. NDA in the DMN. A) Slices showing the thresholded F-statistics map built from the estimated coefficients of the GLM using SPM (http://www.fil.ion.ucl.ac.uk/spm/) overlaid on the T1-weighted image. Top left axial slice: green crosshair locating the center of one NBR region in the Right Lateral Parietal node of the DMN—coinciding with the maximum value of the F-statistics. Bottom left sagittal: green crosshair locating another NBR region in the caudate nucleus. Right slices: Red crosshair locating the center of the PBR region in the right frontal cortex—presumably in the origin of the IEDs. The IEDs were detected using the EEG signal in electrode F8. The approximate location of this electrode is shown with a green circle in the right axial slice to illustrate the possible relation of frontal PBR and the IEDs. The right inset shows a short segment of the preprocessed EEG data where 2/76 IEDs were identified. B) The dark gray curve shows the estimated NN-ARx PBR-HRF and its confidence interval, estimated from the real data around the region marked by the red crosshair in A), and averaged across voxels satisfying *F* ≥ 9.5 within a 7 mm-radius sphere. The light gray curve shows the estimated NN-ARx NBR-HRF and its confidence interval, estimated from the real data around the DMN node marked by the green crosshair in the top left slice in A), and averaged across the voxels satisfying *F* ≥ 70 within a 10 mm-radius sphere. The red and green curves show the unnoisy simulated PBR-HRF and NBR-HRF, respectively, of the fitted EBN model with the estimated value of the recovery time constant: 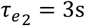.C) For illustration purposes, we also show the temporal behavior of the simulated neuronal activity and BOLD signal in the NBR region (with 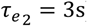), the time series of the real fMRI and the input. To account for the actual relative effect of the IEDs, the amplitude of the input pulses was multiplied by the normalized power of the EEG in F8.

In the second example, a patient with tuberous sclerosis complex (TSC) appears to exhibit ANC just in the edge of a tumor in the left frontal cortex. Although the null hypothesis in the significant voxels could not be rejected with a probability corrected by multiple comparisons, this probability was set to a very low value (*p* < 0.0005) and the minimum cluster size of the significant regions was set to 5 voxels (by decreasing the minimum size of significant voxels, more significant voxels appear in the upper edge of the lesion). Moreover, the HRF were significant according to the permutation test. This HRF corresponded to 50 IEDs identified in electrode F7. Other events were also identified and included as regressors in the linear models. By keeping *ε* constant, the parameter estimation yielded *κ* = 0.51s^−1^.

Figure 10. Possible ANC in the left frontal cortex (cyan crosshair). A) Axial slice of the thresholded F-statistics map built from the estimated coefficients of the GLM using SPM (http://www.fil.ion.ucl.ac.uk/spm/) overlaid on the T1-weighted image. The IEDs were detected using the EEG signal in electrode F7. The approximate location of this electrode is shown with a green circle in the right axial slice to illustrate the possible relation between the NBR region and the IEDs. The right inset shows a short segment of the preprocessed EEG data where 2/50 IEDs were identified. B) The patient suffers from tuberous sclerosis complex. The cyan crosshair—the maximum value of the F-statistics—locates the center of the NBR region, in the perimeter and below one of the patient’s tumors. The tumor is highlighted with the yellow circle in the axial slice and the red arrow in the coronal slice of the T2-weighted image. C) The gray curve shows the estimated NN-ARx HRF and its confidence interval, estimated from the real data, and averaged across the voxels satisfying *F* ≥ 6.5 within a 10 mm-radius sphere with origin in the crosshair in A). The cyan curve shows the unnoisy simulated HRF of the fitted OTT model with the estimated value of the neuro-metabolic coupling gain: *κ* = 0.51s^−1^. D) For illustration purposes, we also show the simulated temporal behavior of *g* and the simulated BOLD in the NBR region (with *κ* = 0.51s^−1^), the time series of the real fMRI and the input. To account for the actual relative effect of the IEDs, the amplitude of the input pulses was multiplied by the normalized power of the EEG in F7.

**Figure 10.**
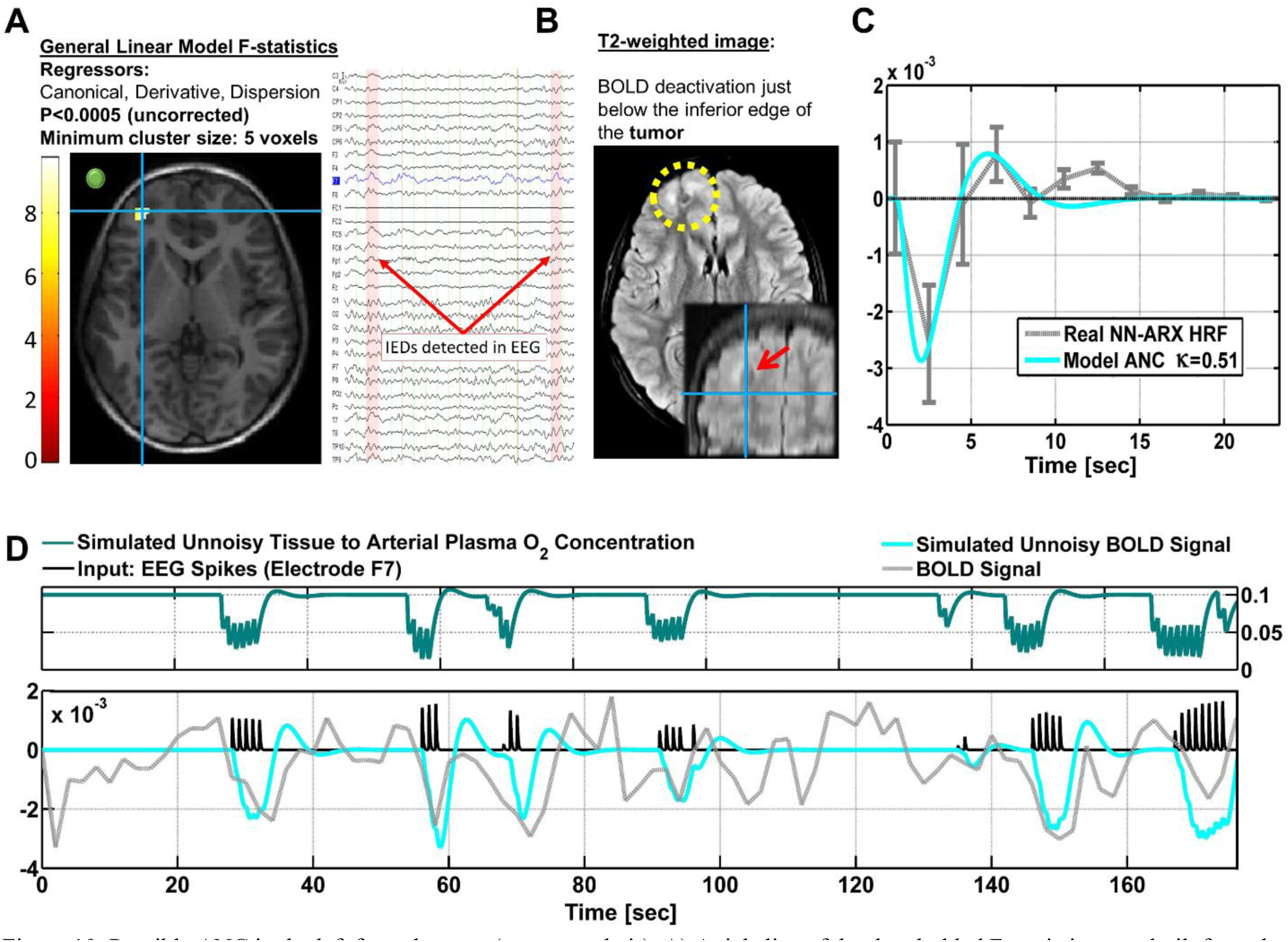
Possible ANC in the left frontal cortex (cyan crosshair). A) Axial slice of the thresholded F-statistics map built from the estimated coefficients of the GLM using SPM (http://www.fil.ion.ucl.ac.uk/spm/) overlaid on the T1-weighted image. The IEDs were detected using the EEG signal in electrode F7. The approximate location of this electrode is shown with a green circle in the right axial slice to illustrate the possible relation between the NBR region and the IEDs. The right inset shows a short segment of the preprocessed EEG data where 2/50 IEDs were identified. B) The patient suffers from tuberous sclerosis complex. The cyan crosshair—the maximum value of the F-statistics—locates the center of the NBR region, in the perimeter and below one of the patient’s tumors. The tumor is highlighted with the yellow circle in the axial slice and the red arrow in the coronal slice of the T2-weighted image. C) The gray curve shows the estimated NN-ARx HRF and its confidence interval, estimated from the real data, and averaged across the voxels satisfying *F ≥ 6.5* within a 10 mm-radius sphere with origin in the crosshair in A). The cyan curve shows the unnoisy simulated HRF of the fitted OTT model with the estimated value of the neuro-metabolic coupling gain: *κ* = 0.51s^−1^. D) For illustration purposes, we also show the simulated temporal behavior of *g* and the simulated BOLD in the NBR region (with *κ* = 0.51s^−1^), the time series of the real fMRI and the input. To account for the actual relative effect of the IEDs, the amplitude of the input pulses was multiplied by the normalized power of the EEG in F7.

Figure 11 shows an IED-related NBR/PBR pair, close to each other, detected in another case. According to the classifier, the mechanism could be lateral inhibition, vascular related, or a combination. The PBR is located in the right superior parietal lobule (SPL) and the NBR region in the right postcentral gyrus (PG), at both sides of the right postcentral sulcus. Assuming ABS, *ρ_A_* = 0.33; while assuming VBB, *ρ_v_* = 0.18. Note that, to detect this NBR/PBR pair, the unsmoothed images had to be used, considerably decreasing statistical significance. This is however the strategy used in (Goense et al., 2012; Harel et al., 2002) to detect close BOLD responses with inverted polarities.

**Figure 11.**
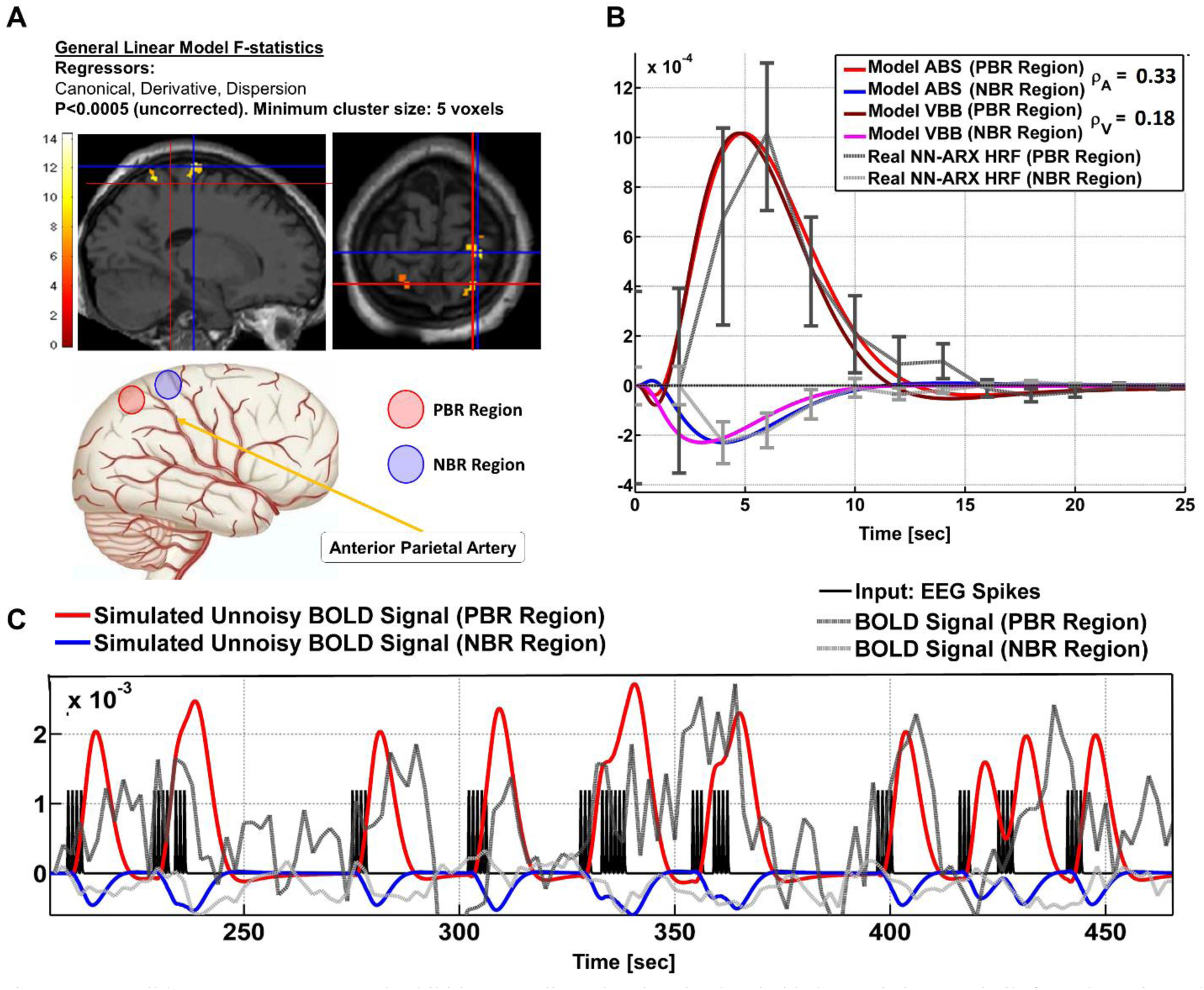
Possible ABS, VBB or Lateral Inhibition. A) Slices showing the thresholded F-statistics map built from the estimated coefficients of the GLM using SPM (http://www.fil.ion.ucl.ac.uk/spm/) overlaid on the T1-weighted image. The blue crosshair locates the center of the NBR region in the Right Postcentral Gyrus; whereas the red crosshair locates the center of the PBR region in the Right Superior Parietal Lobule (SPL), separated by the Postcentral Sulcus (another PBR in the Left SPL was found). The schematic to the right illustrates the ABS mechanism—the regions share the final segment of the Anterior Parietal Artery. B) The dark gray curve shows the estimated NN-ARx PBR-HRF and its confidence interval, estimated from the real data, and averaged across the voxels satisfying *F* ≥ 4.5 within a 10 mm-radius sphere with origin in red crosshair in A). The light gray curve shows the estimated NN-ARx NBR-HRF and its confidence interval, estimated from the real data, and averaged across the voxels satisfying *F* ≥ 5 within a 10 mm-radius sphere with origin in the blue crosshair in A). The light red and blue curves show the unnoisy simulated PBR-HRF, NBR-HRF of the fitted ABS model with the estimated values *ρ*_*A*_ = 0.33. The dark red and magenta curves show the unnoisy simulated PBR-HRF, NBR-HRF of the fitted VBB model with the estimated values *ρ*_*v*_ = 0.18. C) Temporal behavior of the ABS simulated BOLD in the PBR and NBR regions (with the above-mentioned estimated parameters), the time series of the real fMRI and the input used in the NN-ARx estimation.

Figure 11. Possible ABS, VBB or Lateral Inhibition. A) Slices showing the thresholded F-statistics map built from the estimated coefficients of the GLM using SPM (http://www.fil.ion.ucl.ac.uk/spm/) overlaid on the T1-weighted image. The blue crosshair locates the center of the NBR region in the Right Postcentral Gyrus; whereas the red crosshair locates the center of the PBR region in the Right Superior Parietal Lobule (SPL), separated by the Postcentral Sulcus (another PBR in the Left SPL was found). The schematic to the right illustrates the ABS mechanism—the regions share the final segment of the Anterior Parietal Artery. B) The dark gray curve shows the estimated NN-ARx PBR-HRF and its confidence interval, estimated from the real data, and averaged across the voxels satisfying *F*≥4.5 within a 10 mm-radius sphere with origin in red crosshair in A). The light gray curve shows the estimated NN-ARx NBR-HRF and its confidence interval, estimated from the real data, and averaged across the voxels satisfying *F*≥5 within a 10 mm-radius sphere with origin in the blue crosshair in A). The light red and blue curves show the unnoisy simulated PBR-HRF, NBR-HRF of the fitted ABS model with the estimated values *ρ_A_*=0.33. The dark red and magenta curves show the unnoisy simulated PBR-HRF, NBR-HRF of the fitted VBB model with the estimated values *ρ_v_*=0.18. C) Temporal behavior of the ABS simulated BOLD in the PBR and NBR regions (with the above-mentioned estimated parameters), the time series of the real fMRI and the input used in the NN-ARx estimation.

## Discussion

We have provided a theoretical framework to study the mechanisms that generate negative BOLD responses (NBR) during event related or block design fMRI experiments. According to our biophysical models, these NBRs depend on some crucial parameters that determine their amplitude and shape. These parameters are determined by the anatomical, functional and physiological characteristics of the brain or the tissue involved in the generation of the responses.

Inhibition-related phenomena, i.e. LCI and NDA, require accounting for the imbalance between the activation of inhibitory and excitatory neuronal states and their respective connectivities. Note that the two state model was already used by (Havlicek et al., 2017, 2015) to explain NBR during static and flickering visual stimulation. With this approach, it is possible to evaluate the competing contributions to BOLD responses of both activation and deactivation of neuronal activity. In this paper, we proposed a simplified quantitative framework where we analyze the combination of neuronal connectivity which yield PBRs and N. Contrary to (Havlicek et al., 2017, 2015), our two-state model not only accounts for an input to the excitatory state, but also to the inhibitory one. We believe that this inclusion is appropriate since the inputs can be seen as either thalamocortical/corticocortical inputs to neurons in the granular layer, relayed to both excitatory and inhibitory populations, or modulation of firing thresholds or conductance in both populations. Significant NBRs cannot be obtained unless inputs are affecting the inhibitory state. In remote-LCI this can be seen as the effect of collaterals in a lateral inhibition network (Priebe and Ferster, 2008; Shamma, 1985) or callosal inhibition (Bloom and Hynd, 2005). In the case of local-LCI (which is actually another type of lateral inhibition) we turned our attention to the formation of apparently contradicting NBR by local neuronal activations, such as IEDs in epilepsy (Pittau et al., 2013). It is believed that IEDs are generated by a brief increase in excitatory feedback gains and decrease in firing thresholds (Ayala et al., 1973). The latter is compatible with the view of a short input to both neuronal states. But only the balance of inhibitory-excitatory connectivities can determine which polarity of BOLD response prevails. Note that the “wave” component of epileptic events is believed to be related to a decrease of potassium conductance (or dis-facilitation of potassium currents) (Neckelmann et al., 2000) or robust hyperpolarization in III/V layer pyramidal cells (Pollen, 1964). Thus, a decrease of synaptic gains in the excitatory population exceeding those of the inhibitory population, i.e. 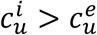 and *c*_*i*_ > *c*_*e*_, will provide the necessary conditions for an NBR to occur. Assuming *c*_*i*_ > *c*_*e*_ is actually compatible with inhibitory synapses closer to the soma (Megi´as et al., 2001; Villa and Nedivi, 2016) and it has been adopted in the computational neuroscience literature (David et al., 2006; Nunez, 1995; Robinson, 2005). The spike (positive) and slow wave (negative) change in neuronal activity shown in the simulations of Figure 2 is apparently in contradiction with the EEG traces observed during these events, in which voltage deflections for both phases have the same polarity (Pittau et al., 2013). Note however that neuronal activity does not directly map to EEG (Riera et al., 2006). The behavior of *n* is very similar to *n*_*e*_ (not shown). The latter variable might be regarded as mostly representing the activity of pyramidal cells. We expect that a depolarization (excitation) of these cells usually occurs as a result of synapses in the apical dendrites forming a current sink with its corresponding source in deeper layers. On the contrary, hyperpolarization (inhibition) mostly occurs near the basal dendrites and soma, generating a current source with its corresponding sink in more superficial layers. In both situations a current dipole along the basal-to-apical direction is formed, yielding an EEG trace with the same polarity. It is important to clarify however that the states do not necessarily represent the synaptic activity of lumped populations of excitatory and inhibitory neurons, but competing states of self-organizing synaptic activity determined by several mechanisms such as adaptation, gain-control, refractoriness, polysynaptic connectivity from recurrent axonal collaterals, among others. (Marreiros et al., 2008). Although beyond the scope of this paper, a more realistic approach which includes neural masses with linear dendritic transfer functions and nonlinear membrane potential to firing rates transformations (David et al., 2006; Sotero et al., 2007), adaptation and facilitation (Robinson, 2011), and subcortical structures (van Albada and Robinson, 2009) has to be considered in the future.

The nature of the mechanisms described in this paper determines which time constants should be assumed for the states. In NDA, the recovery of the neuronal activity of the disrupted network is characterized by a large 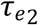 since it actually depends on the way the different nodes conflictedly interact to effectively “shut down” and recover the network when a strong stimulus affects one of its nodes. The way process occurs is unknown. Note that brain dynamics operate near criticality (Deco et al., 2011; Deco and Jirsa, 2012; Ghosh et al., 2008), i.e. on the brink to instability. Neuronal activities in this situation require higher recovery time to reach equilibria after stimulation, and are associated with large-scale dependencies and scale invariance (Haken, 1983). On the other hand, the center manifold theorem states that a high-dimensional system in a subcritical state can be projected to a lower-dimensional manifold, i.e. a subset of networks, when the system is stimulated. This is compatible with the appearance of networks, such as the DMN, that outlasts the stimulation. To study such dynamics in earnest, multiple neuronal populations must be modeled at several nodes of an embedded anatomical network, and the sensitivity of the functional network integrity will depend on the type of node that is affected by the excitatory external input (Spiegler et al., 2016). Examples of this type of modeling are (Jirsa et al., 2017), where the effect of epileptic activity and interventions are investigated using a mathematical model of an epileptic brain, and (Kunze et al., 2016), where the effect of transcranial direct current stimulation on functional connectivity is studied. Such characterization is beyond the scope of this paper. Our goal is to provide a simpler model for the neuronal dynamics that determine the NBR waveform to investigate how it differentiates from the rest of the mechanisms.

If neurovascular coupling is kept constant, the parameter which is highly correlated with NBR amplitude will be *κ*. A disproportionate increase of this value leads to ANC. In this mechanism, the NBR can be seen as an exaggerated initial dip. This phenomenon cannot be observed using linear methods to deconvolve the HRF when the stimulation is presented for several seconds at a high frequency. This happens because sustained neuronal activity with excessively high coupling between the activity and the CMRO_2_ depletes the O_2_ in the tissue until the end of the stimulation. Without O_2_ in the tissue the total blood O_2_ concentration returns to its baseline since the CBF is constant during the sustained stimulation; and the oxygen extraction operates according to the oxygen limitation regime (Buxton and Frank, 1997) in which the extraction is lower than the baseline, the relative dHB content is consequently lower in the venules and the response becomes positive during the rest of the stimulation.

To explain the two vascular phenomena, i.e. ABS and VBB, we coupled two windkessels by an artery or a vein, respectively. Simulations indicate that the parameter which determines the NBR amplitude is the resistance of the vessel (artery or vein), relative to the total steady state resistance of the vasculature within the tissue, i.e. the arterioles, capillaries and venules. Note that, VAN (Boas et al., 2008) predicts a negativity in CBF and O2 saturation surrounding a positivity of the same magnitudes. More closely, the surround CBV decreased in the arterioles and increased in the venules, suggesting reduced pressure in the former and backpressure in the latter. In that model, each parent arteriole is divided into two children; and two parent venules merge in a child. Besides, since it is a model of a circuit of multiple vascular compartments, the vessels need to be modeled with blood resistance. Therefore, both ABS and VBB mechanisms are present. On the other hand, compared to the response reported in (Harel et al., 2002) for 20s of stimulation^1^, the simulated NBR using the ABS sub-model also lacks a post-stimulus overshoot. That paper reports a CBV response that peaks and prolongs after the cessation of the stimulus, contrary to our simulations, which however resemble the CBV response reported in (Ma et al., 2017)^2^, presumably for the same type of phenomena. The NBR obtained using our VBB model for 20-second stimulation lacks the post-stimulus overshoot reported by (Goense et al., 2012)^3^. Authors suggested that the NBR might be the result of the combination of blood backpressure and neuronal inhibition (Bandettini, 2012; Goense et al., 2012). That would explain the presence of the post-stimulus overshoot, which is in contrast to our pure vascular simulations. This undershoot is shown to be determined by the dynamics of the inhibitory neuronal state (Havlicek et al., 2017). Other causes including vein delayed compliance might also explain these transient.

In this paper, we focus in the possibility of detecting and classifying the NBR mechanisms using the HRFs extracted from BOLD fMRI signals. We use linear models for the detection and estimation of the HRFs, i.e. GLM and NN-ARx, respectively. Although linear models have been used to detect and reconstruct BOLD responses for decades, even before addressing the nonlinear characteristics of BOLD signals (Buxton et al., 2004; Friston, 2002), we decided to analyze if linear models are applicable for our data. We found that the NDA and ANC are significantly nonlinear. We proposed a correction method based on a calibration of the amplitude of the fMRI signals by multiplying it by the frequency profile of the input, calculated using a moving average window. More rigorous approaches are beyond the scope of this paper. Future work must consider the use of wavelet-based time-frequency analysis methods (Wacker and Witte, 2013) to build the calibration profile or directly deal with the nonlinearities in the regression by using generalized nonlinear models and nonlinear autoregressive models (Seber and Wild, 1989). Additionally, we can also increase detection sensitivity by including higher order Volterra kernels in the GLM (Friston, 2002).

We investigated if the NBR mechanisms can be solely classified from their BOLD responses. This is important for clinical applications when only standard fMRI paradigms are available or designed. The prediction costs in Figure 7 shows that the machine learning classifier is unable to differentiate LCI, ABS and VBB in almost 30% of the cases. In interictal epilepsy, this might be worse since there is a loss of sensitivity and specificity related to the usual misclassifications of IEDs, which is expert dependent. These mechanisms are different in nature and our model predicts different responses when using other imaging or recording modalities. At the expense of experimental feasibility, MION (Harel et al., 2002) and/or VASO (Goense et al., 2012) can be used to measure CBV concurrent with BOLD signals. Measurements of neuronal activity can be incorporated using LFP and MUA (Schridde et al., 2008) or EEG (Maggioni et al., 2016). In addition, CBF can be also included using Arterial Spin Labeling (ASL) (Shmuel et al., 2002) and/or FAIR (Goense et al., 2012). With these multimodal observations, we foresee a significant increase in the margins of classification of the NBR mechanisms, even in the case more than one is present in the same region (Goense et al., 2012; Shmuel et al., 2006). In that respect, we point out that we decided to study particular cases of our model for simplicity, numerical stability and parsimony. Future work is needed to cope with the classification and identification of a more realistic “continuum” of multi-mechanistic phenomena.

We explored the existence of the IED-related NBR mechanisms in a set of patients with focal epilepsy. In the first example, a clear NDA was detected in the DMN. These DMN deactivations have been systematically reported in the literature for temporal lobe epilepsy (TLE) (Kobayashi et al., 2006; Laufs et al., 2007) and for other types of focal epilepsy (Fahoum et al., 2013, 2012). The pattern of deactivation depicted in Figure 9, with a predominance in the parietal node of the DMN ipsilateral to the focus, is similar to that reported in (Fahoum et al., 2013) for four out of five patients, with concomitant decrease of electrophysiological activity. Our results suggest that the PBR region might have afferents on the anterior part of the caudate nucleus that relay to central nodes of the DMN. This is consistent with the hypothesis of widespread secondary inhibition of non-seizing cortical regions via basal ganglia (Norden and Blumenfeld, 2002). The temporal profile of the PBR HRF in the right IFG experienced an unpredicted decay (or rebound) correlated with the amplitude of the NBR in the DMN nodes. This might be seen as an interruption of the PBR mechanism by inhibitory afferents coming from the regions exhibiting NDA, which in this case are present all around the right IFG. This might have implications in the interpretations of BOLD responses during IEDs or stimulation paradigms. If the location of the PBR is close to an affected RSN node, its HRF waveform might be misleading of the actual underlying PBR mechanism, due to either the interaction between mechanisms, i.e. inhibitory inputs from the NBR to the PBR region, or the effect of the BOLD spatial point spread function. This rebound can be also explained as an increase in dHb due to an increase in neuronal activity in the NBR region, or even a vascular reallocation phenomena, as suggested by (Hu and Huang, 2015). They observed positive and negative optical responses, concurrent with LFP and MUA measurements, in rats during hindlimb electrical stimulation. Finally, it has been hypothesized that NDA is a disruption of RSN provoking a reduction of consciousness and cognitive reserve (Fahoum et al., 2013). Interestingly, our results suggest that a recovery from this disrupted state is not instantaneous. In our data, the estimated value for the recovery time constant was 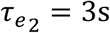.

The NBR in our second case was classified as ANC. This mechanism is related to a decrease in the CBF/CMRO_2_ balance. This was the hypothesis to explain NBRs in the hippocampus of rats during bicuculline-induced generalized tonic-clonic seizures (Schridde et al., 2008). They reported that, even with higher LFP/MUA activity in the hippocampus, as compared to the cortex, the CBF was lower and the CMRO_2_ was higher, yielding to NBRs in the hippocampus. Quantitatively, the unbalance corresponds to a decrease in the ratio *ε*/*κ* in the OTT model (Zheng et al., 2002), which is around 0.40.05/ = 8 for normal positive responses. Confirmation of the results in (Schridde et al., 2008) was provided by (Song et al., 2016), who estimated this ratio about 5-fold smaller in rat with focal cortical seizures, mainly owing to an increase in *κ*. Although the rCBF/CMRO_2_ coupling was reported to be preserved in human IEDs without any apparent lesion (Stefanovic et al., 2005), we do not discard the possibility of an unbalance produced by a more critical state of tissue pathology. For example, the typical calcification of the surrounding blood vessels present in TSC tumors (Gallagher et al., 2010) could hamper the expected IED-induced increase of rCBF. We estimated *κ* = 0.51s^−1^ for the IED-related ANC mechanism around the lesion, which yields 0.280.51/ = 0.55, 14 times smaller than the normal values.

In our last example, we find an NBR in the Superior Parietal Lobule (SPL) and a PBR in the Postcentral Gyrus (PG). The NBR classified as either lateral inhibition (LI) which would occur via either U-fibers or vascular phenomena. However, we cast doubt on LI since we believe an inhibitory pathway from SPL to PG is rather weak. Note that the PG hosts the primary somatosensory cortex (S1) (Geyer et al., 2000), which is a granular cortex that mainly receives somatotopic feedforward afferents from the Ventral Posterolateral and Posteromedial relay nuclei (VPL and VPM) of the thalamus (Cappe et al., 2011). In addition, the SPL, involved in transforming visual information in complex motor planning, has efferent pathways mainly to the premotor supplementary motor cortices in the precentral gyrus. A top-down inhibition from higher areas to the somatosensory area is mainly via efferents in the prefrontal cortex. Moreover, both DSI-based connectivity (Hagmann et al., 2007) and cortical thickness based connectivity (Joshi et al., 2010) between SPL and PG are rather low. The mechanism might be ABS. Note that the detected BOLD responses are in the vascular domain of the middle central artery, at both sides of the postcentral sulcus. Thus, they might be sharing a final segment of the anterior parietal artery. Although we do not discard VBB, in which the regions could be sharing some anastomic vein feeding the central sulcal vein or a branch of the superior anastomic vein of Trolard, it has been reported that venous CBV changes are only relevant for longer stimuli (Uludağ and Blinder, 2018).

The proper identification of these mechanisms has important clinical implications in epilepsy. For example, we believe that only the PBR and NBR that occurs in the seizure onset zones (SOZ) are clinically relevant—excluding the NBR regions associated to NDA or vascular phenomena. In normal stimulation paradigms, such as event related and block designs, one must be cautious in the interpretation of NBRs as neuronal inhibition, disruption of certain brain states or vascular phenomena without a direct functional interpretation.

It is worth noting the foreseeable boost that BOLD modeling will have with the advent of new and optimized sequences in high field spin-echo fMRI, with the considerably improved ability to measure high resolution layer-dependent BOLD images and correlates of rCBV and rCBF (Bandettini, 2014; Feinberg and Yacoub, 2012; Goense et al., 2016, 2012; Harel et al., 2010; Huber et al., 2015, 2014; Polimeni and Uludağ, 2018). This allows for the construction and estimation of more detailed models of BOLD generation, through understanding of the actual role of arteries, capillaries and veins in the generation of these observables and the possible biases that the variability of neuro-vascular/metabolic coupling, CBV and SNR across layers could introduce. For example, it has been reported that the baseline CBV distribution varies over cortical layers biasing fMRI signal to layers with high CBV values (Uludağ and Blinder, 2017). This affects the interpretation of what the contribution of the different vascular compartments to the average low-resolution BOLD response is.

In this paper, we discussed several types of NBR mechanisms that are related to the balance between excitatory and inhibitory neuronal states, neurovascular and neurometabolic couplings and blood properties. These were accommodated as particular cases of a common biophysical model. It must be mentioned that there are other types of NBR mechanism that are not explained by our model. For example, a study demonstrates that noxious electrical stimulation induces CBV decreases, underlying NBR, in the Caudate-Putamen which is associated with increased neural activity (Shih et al., 2009). It is believed that activated neurons respond directly to pain modulation while endogenous neurotransmission (dopamine) induces a low energy tone CBV decrease to maintain the quiescent neural activity (Shih et al., 2009). Another example is NBR observed due to volume changes of the CSF adjacent to the lateral ventricle of macaque V1 (Goense et al., 2016) and in periventricular CSF and large veins (close to activations areas) in humans (Bianciardi et al., 2011), due to a mismatch between CBV increase and oxygenation changes in regions devoid of tissue. Another NBR not explained by our model is observed in white matter due to vasoconstrictive events and long blood transit times in the medullary vasculature of this tissue, delaying the blood oxygen changes relative to CBV effects (Özbay et al., 2018).

## Conclusions

In summary, five NBR mechanisms were identified and quantitatively described under the same theoretical framework. The NBR can be detected in realistic fMRI scans and some of them can be classified according to their HRF. For lateral and contralateral inhibition and vascular mechanisms, complementary modalities of data are needed. We found examples of some of these mechanisms in real epileptic EEG-fMRI data and estimated the parameters that are determinant in the formation of the NBR mechanisms.

## Acknowledgments

We appreciate the members of the Neuronal Dynamic Mass Laboratory, Prof. Pedro A. Valdes-Sosa and Prof. Louis Lemeiux for their useful comments and suggestions during the preparation of this manuscript.

## Funding sources

This work was supported by the National Institutes of Health (R56NS094784-01A1).

## Competing interests statement

The authors declare no conflict of interest

## Appendix. Derivation of Stealing/Backpressure models

**Figure 12.**
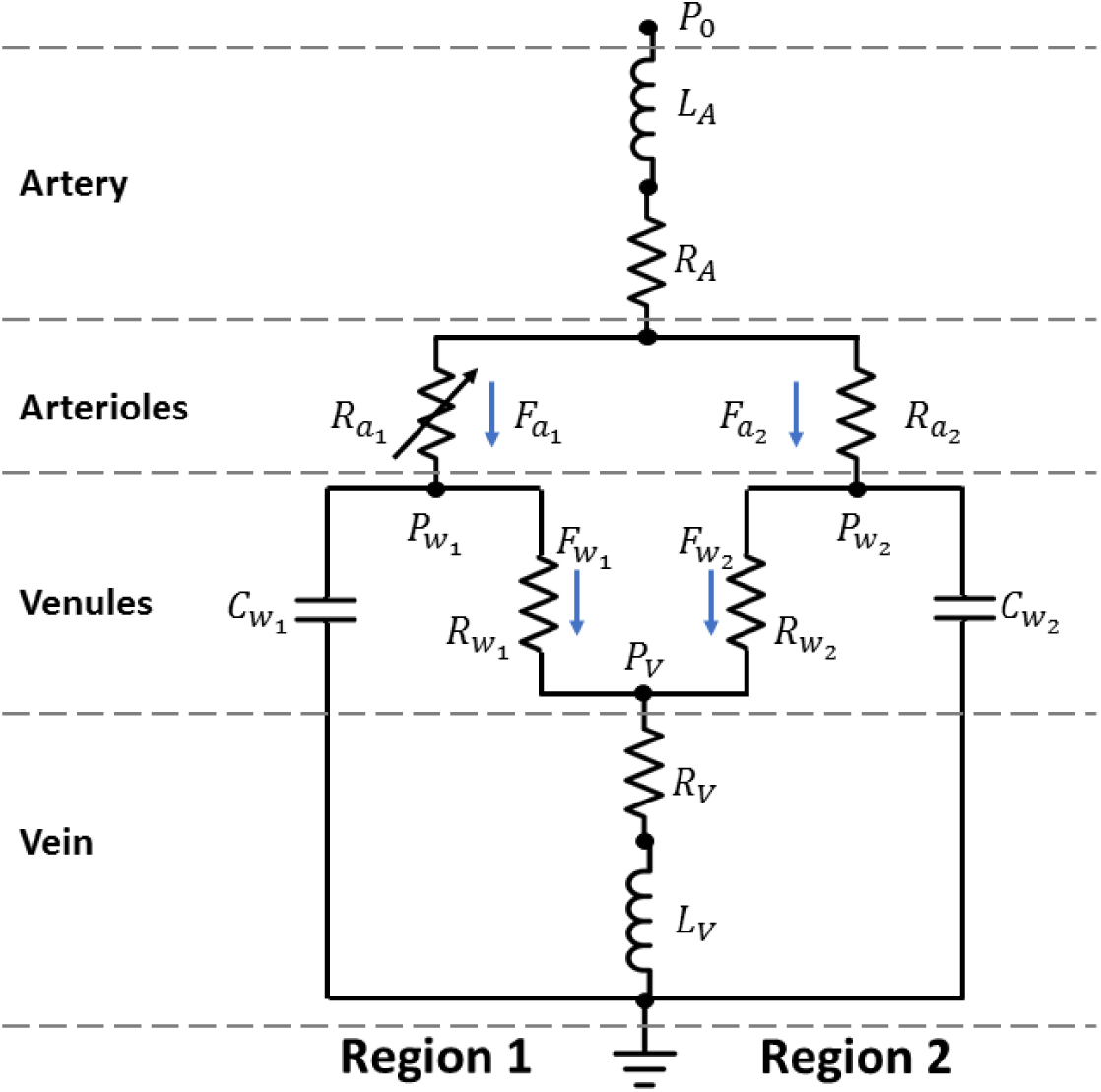
Figure 12. Circuit equivalent to the vascular part of the model in Figure 1. Flow (*F*) and pressure (*P*) amount current and pressure, respectively. The arteriole resistance, controlled by the vasoactive signal, is variable. Both regions are connected by a common supplying artery and draining vein, characterized by blood resistance and inertia (inductive element). The windkessels are characterized by nonlinear resistance (*R*_*w*_) and capacitance/compliance (*C*_*w*_). The circuit is under a constant cerebral blood pressure (voltage, *P*_0_).

In this appendix we derive equations (7), (8) and (9) of the main text. The vascular part of the model in Figure 1 is equivalent to the circuit in Figure 12. To ease the understanding of the following derivations, we define the following mathematical structures:

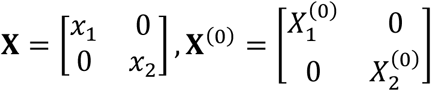

According to the first Kirchhoff’s law, the change in the volume (i.e., the capacitive current) of the *k*^*th*^ windkessel is given by:

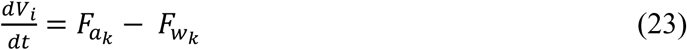

Dividing equation (23) by 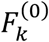, where 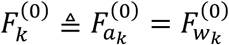, and considering the central volume principle, i.e. 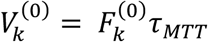, we obtain equation (7):

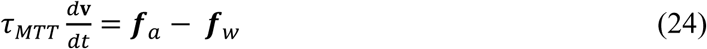

Second Kirchhoff’s law along the *k*^*th*^ windkessel yields the following system of differential equations:

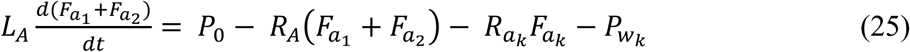

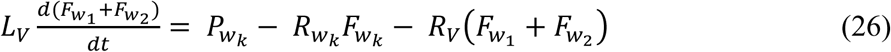

Windkessel theory for diminished reserved volume and nonlinear compliances (Mandeville 1998) for an arbitrary state variable *A*_*k*_ states:

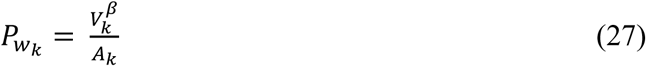

Using equation (27) and defining *L*_*A*_ ≜ *τ_A_R_A_* and *L*_*v*_ ≜ *τ_v_R_v_*, equations (25) and (26) can be written using matrix notation as follows:

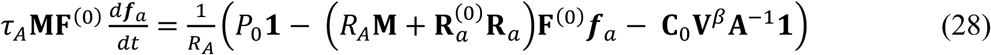

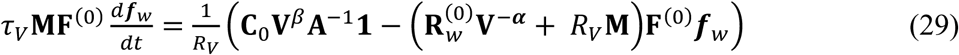

Where 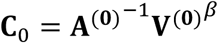 and M=11^T^.

At baseline, equation (29) simplifies and yields 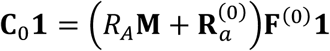. Using this, the steady state form of equation (29) yields:

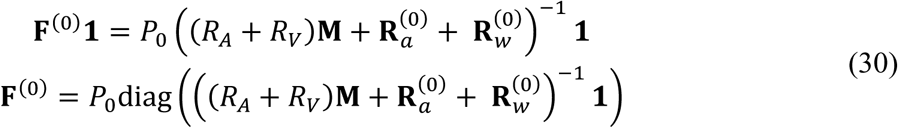

If we substitute **F**^(0)^ in (30) we realize neither this equation nor equation (29) depend on *P*_0_. Thus, we can set *P*_0_ = 1 without loss of generality. On the other hand, we can choose a combination of resistances and baseline **F**^(0)^ that satisfies equation (15). Therefore, we can set a “gauge”, say **1**^T^**F**^(0)^**1** = 2, which implies

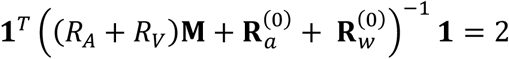

In this paper, the resistance configuration is identical for both regions, i.e. 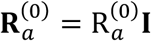 and 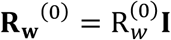. This implies:

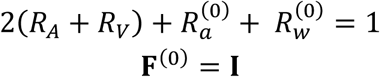

Therefore, the final set of differential equations for the arteriole and venule normalized flows are:

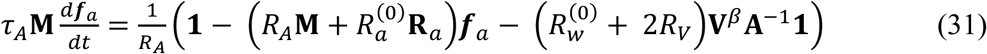

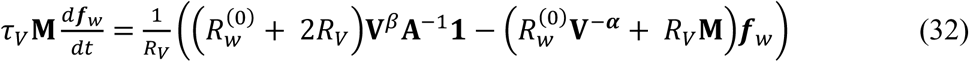

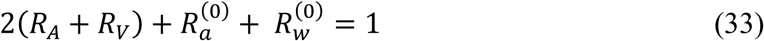

Let set *R*_*A*_ = *R*_*v*_ = 0. This corresponds to a very high diameter of the vessel. Note also that these resistance decay much faster than the increase of the inertial times with increasing diameter (see (19) and (20)), thus the left sides of (31) and (32) become zero. The simplified versions of these equations, after substituting in (24) and using (27), yield (Mandeville et al., 1999) uncoupled equations:

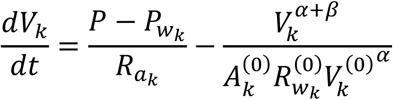

We adopted the definition of (Zheng and Mayhew, 2009) for the variable *A*_*k*_. To be fully compatible with (Mandeville et al., 1999), we must change *A*_*k*_ to 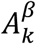

## Footnotes

1 See Figure 3 of that paper

2 See Figure 2 of that paper

3 See Figure 3 of that paper

